# Instability of aquaglyceroporin (AQP) 2 contributes to drug resistance in *Trypanosoma brucei*

**DOI:** 10.1101/2020.02.22.960963

**Authors:** Juan F. Quintana, Juan Bueren-Calabuig, Fabio Zuccotto, Harry P. de Koning, David Horn, Mark C. Field

## Abstract

Defining mode of action is vital for both developing new drugs and predicting potential resistance mechanisms. African trypanosome pentamidine and melarsoprol sensitivity is predominantly mediated by aquaglyceroporin 2 (TbAQP2), a channel associated with water/glycerol transport. TbAQP2 is expressed at the flagellar pocket membrane and chimerisation with TbAQP3 renders parasites resistant to both drugs. Two models for how TbAQP2 mediates pentamidine sensitivity have emerged; that TbAQP2 mediates pentamidine translocation or via binding to TbAQP2, with subsequent endocytosis, but trafficking and regulation of TbAQPs is uncharacterised. We demonstrate that TbAQP2 is organised as a high order complex, is ubiquitylated and transported to the lysosome. Unexpectedly, mutation of potential ubiquitin conjugation sites, i.e. cytoplasmic lysine residues, reduced folding and tetramerization efficiency and triggered ER retention. Moreover, TbAQP2/TbAQP3 chimerisation also leads to impaired oligomerisation, mislocalisation, and increased turnover. These data suggest that TbAQP2 stability is highly sensitive to mutation and contributes towards emergence of drug resistance.

## Introduction

Human African trypanosomiasis (HAT) is a neglected tropical disease affecting sub-Saharan countries [1–4]. HAT progresses by two stages: a haemolymphatic stage, in which the parasite successfully colonises the bloodstream, lymphatics, skin, adipose tissue and organs and a meningoencephalic stage characterised by the emergence of parasites in the central nervous system (CNS) [2,5]. Several drugs are used to treat HAT; currently suramin and pentamidine are the drugs of choice for treatment of the haemolymphatic stage of *T. brucei rhodesiense* and *T. brucei gambiense* infections respectively, whereas melarsoprol, eflornithine or combined nifurtimox-eflornithine (NECT) therapy are recommended for the meningoencephalic stage [6,7].

Two new drugs, fexinidazole and acoziborole, recently completed clinical trials and opened a new front in HAT chemotherapy [8,9]. Drug development, successful public health initiatives and active case-monitoring programs have all contributed to the anticipated eradication of *gambiense* HAT as a major public health problem in the coming decade [10]. However, vigilance and understanding of drug mechanisms and possible resistance pathways remain essential to maintain this situation, and *rhodesiense* HAT cannot be eliminated in this way as it is highly zoonotic [11]. Genome-wide RNAi screens identified a number of genes associated with pentamidine sensitivity that, together with evidence from melarsoprol-pentamidine cross-resistance (MPXR), identified aquaglyceroporin 2 as the primary determinant for drug-uptake [12,13], alongside lesser roles for the TbAT1/P2 aminopurine transporter and the Low Affinity Pentamidine transporter LAPT1 [14].

Aquaglyceroporins (AQPs) are an ancient family of multi-pass membrane proteins, containing both aquaporins that exclusively transport water and aquaglyceroporins that transport both water and uncharged low molecular weight solutes [15–17]. The *T. brucei* genome encodes three AQPs (TbAQP1-3) [18], all of which are nonessential but do control osmoregulation and glycerol transport [13,19– 24]. TbAQP1 (Tb927.6.1520) and TbAQP2 (Tb927.10.14170) are typically localised to the flagellum and flagellar pocket respectively, whereas TbAQP3 (Tb927.10.14160) is associated with bulk plasma membrane [13,21,23]. TbAQP2 and TbAQP3 are the product of a recent gene duplication within the African trypanosome lineage [13,21,23,25].

A selectivity filter restricts the size and properties of solutes that can effectively pass through the AQP pore [15–17]. In TbAQP1 and TbAQP3, this is formed by two constrictions of the channel: the canonical “NPA” within two half alpha helices and a narrower “aromatic/Arginine” (ar/R) motif (Fig. 1A) [13,25,26]. Significantly, TbAQP2 does not retain the canonical configuration but displays an unconventional “NPS/NSA” cation filter motif. Similarly, the ar/R motif is replaced by a neutral leucine at position 264 (L264), followed by aliphatic rather than aromatic residues (A88, I110, V249 and L258), which are equivalent to the “IVLL” motif observed in the selectivity pore of other AQPs [13,23]. These substitutions may permit TbAQP2 to transport larger solutes, including pentamidine (340 Da) [23]. However, pentamidine also binds TbAQP2 with nanomolar affinity and replacement of the Leucine 264 by arginine abolishes binding, leading to resistance [22], consistent with a proposed hypothesis that pentamidine sensitivity might be mediated by high affinity binding of pentamidine to TbAQP2 and internalisation *via* endocytosis [22]. It is also plausible that pentamidine exploits both channel activity and endocytosis of TbAQP2 to gain entry to the trypanosome cytoplasm.

**Figure 1.**
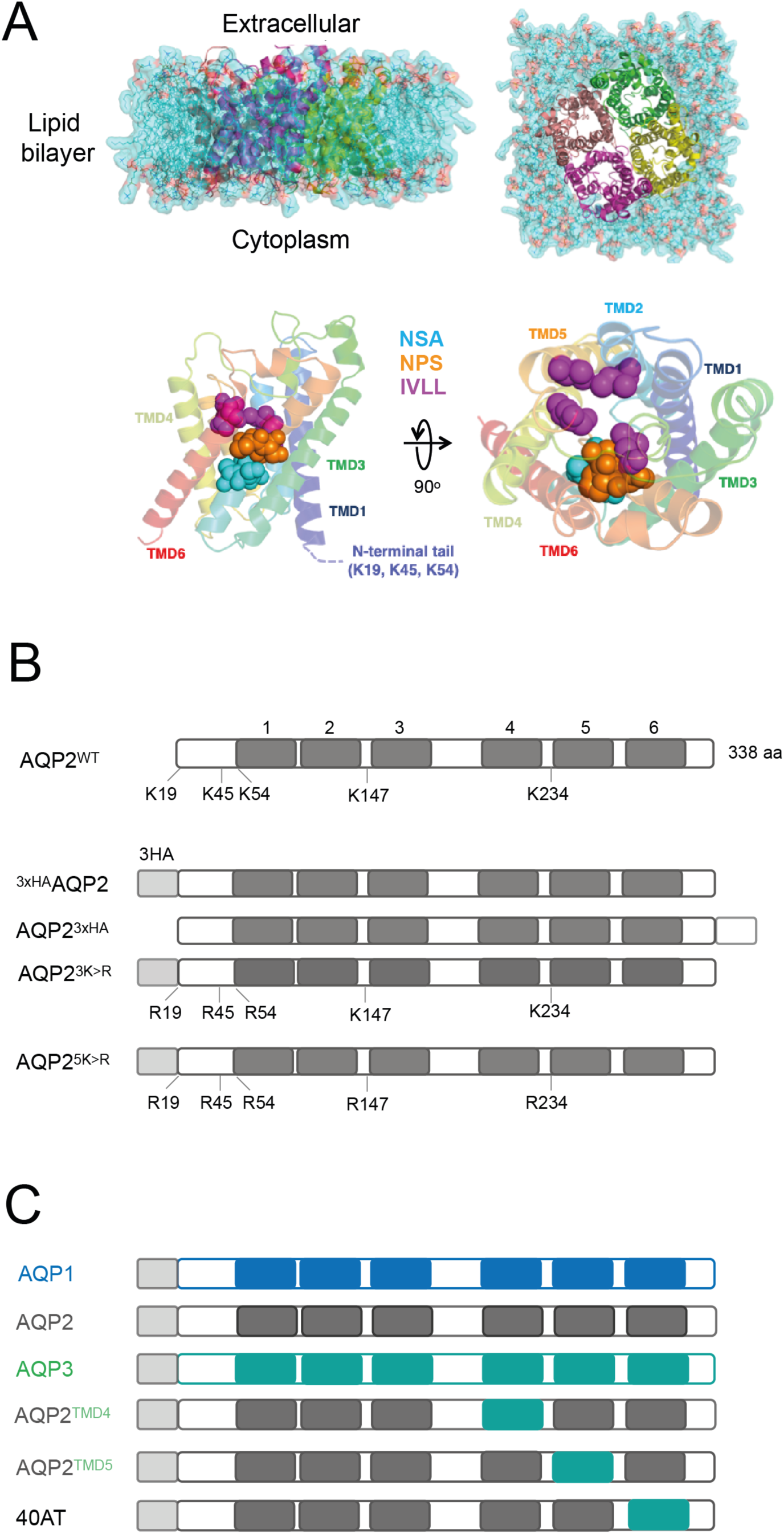
Schematic representation of constructs used in this study. **A)** 3D structural predictions of the AQP2 harbouring three haemagglutinin tags at either terminus. **Top panel**; lateral and cytoplasmic face view of simulated model of *T. brucei* AQP2 tetramer embedded in a POPC lipid bilayer. Lipids are shown in surface and line representations in cyan. Each monomer of AQP2 is shown in cartoon representation. **Bottom panel**; lateral and cytoplasmic face view of *T. brucei* AQP2 showing key amino acids (in spheres) from NSA (cyan), NPS (orange) and IVLL (magenta) domains. **B)** N- and C-terminal tagged TbAQP2 variants with a tandem of three hemagglutinin (3xHA) epitopes. Positions of predicted *trans*-membrane domains (TMD) are indicated with numbers above solid blocks. Similarly, lysine residues that were manipulated in this study are highlighted. **C)** Wild type TbAQP1 (blue), TbAQP2 (grey), TbAQP3 (green), and chimeras used (40AT, AQP2^TMD4^, and AQP2^TMD5^). TMDs for AQP1, 2 and 3 are shown as blocks and in blue, grey and green, respectively.

Melarsoprol-pentamidine cross-resistant strains and field isolates from relapse patients all possess mutations at the locus encoding TbAQP2, including deletions, single nucleotide polymorphisms and chimerisation [27–30]. Pentamidine-resistant trypanosomes from a cohort of relapse patients from the Democratic Republic of Congo also have TbAQP2 chimerisation with the coding sequence for the C-terminal *trans-*membrane domain replaced by TbAQP3 and in most cases without altering the selectivity filter characteristic of TbAQP2 (NSA/NPS – IVLL motifs) [29,31,32]. These observations indicate that, in addition to the sequence at the selectivity pore, other features are likely to impact drug uptake and transport in *T. brucei* [33].

Here, we set out to investigate TbAQP2 trafficking and to understand the basis of drug resistance in chimeras where the selectivity filter remains intact. We find that AQPs form a tetramer of tetramers (4×4) quaternary structure which correlates with high stability, flagellar pocket localisation and functionality. Furthermore, we demonstrate that TbAQP2 is ubiquitylated and highly sensitive to mutation of cytoplasmically oriented lysine residues. Finally, we find that chimerisation of TbAQP2, as observed in DRC patients, leads to protein instability and mislocalisation, thus explaining the basis for drug resistance in clinical isolates of *T. brucei*.

## Materials and methods

### Cell culture and drug sensitivity

Bloodstream form (BSF) *T. brucei* 2T1 and all derivatives were cultured in HMI-11 (supplemented with 10% heat-inactivated fetal bovine serum (FBS), 100 U/ml penicillin, 100 U/ml streptomycin and 2 mM L-glutamine) at 37°C with 5% CO_2_ in a humid atmosphere in non-adherent culture flasks with vented caps at densities between 1 x 10^5^ and 1.5 x 10^6^ cells/ml. 2T1 cells were maintained in the presence of phleomycin (1 μg/ml) and puromycin (1 μg/ml). Following transformation, cells were selected and maintained with hygromycin (2.5 μg/ml) or phleomycin (1 μg/ml) as appropriate. EC_50_ determinations were performed using AlamarBlue® (resazurin sodium salt) as described [34,35], with 5 mM glycerol added as appropriate; drug exposure was for 66 hours and AlamarBlue® incubation overnight. Plates were read on an Infinite 200Pro plate-reader (Tecan) with the following parameters: excitation, 530nm; emission 585 nm; filter cut-off, 570 nm. Proliferation was monitored by dilution to 1 x 10^5^ cells/ml and counting daily. For transfections, 3 x 10^7^ bloodstream form cells were harvested by centrifugation and transfected with 5-10 μg of linearized plasmid DNA using an Amaxa Nucleofector II (Lonza) with program X-001. Bafilomycin A1, MG132, Salicylhydroxamic acid (SHAM), glycerol, AlamarBlue®, pentamidine and ammonium chloride were all from Sigma.

### Recombinant DNA manipulation

To express N- or C-terminal HA-tagged AQP constructs, a tandem of three HA tags was inserted by PCR using the primers: Tb^3xHA^_AQP2_Fwd (HindIII): CCCAAGCTTGGGATGTACCCATACGATGTTCCAGATTACGCTTACCCATACGATGTTCCAGATTACGCTTACCCATACGATGTTCCAGATTACGCTCAGAGCCAACCAGACAATGTG and Tb^3xHA^_AQP2_Rev (BamHI): CGCGGATCCGCGTTAGTGTGGAAGAAAATATTTGTAC. The PCR products were inserted into pRPa^TAG^ [12] after digestion with BamHI/HindIII. All constructs were verified by sequencing (MRC-PPU DNA Sequencing facility, University of Dundee). Prior to introduction into trypanosomes pRPa^TAG^ constructs were linearized with AscI and purified/sterilized by phenol:chloroform extraction. TbAQP2, with all lysine residues predicted facing the cytoplasm mutated (TbAQP2^5K>R^) was designed and synthesized by GenScript and verified by sequencing. Point mutations rescuing individual lysine residues were introduced using the Q5 Site-Directed Mutagenesis Kit (NEB) and confirmed by sequencing. Tagging of lysine mutants was conducted as above.

### Imaging

Antibodies were used at the following dilutions: rat anti-HA IgG_1_ (clone 3F10; Sigma) at 1:1000, rabbit anti-ISG75 (in house) at 1:500, mouse anti-p67 (from J. Bangs) at 1:500. Secondary antibodies (Thermo) were at: anti-rat Alexa-568 at 1:1,000, anti-rabbit Alexa-488 at 1: 1,000, anti-mouse Alexa-488 at 1: 1,000. Coverslips were mounted using Vectashield mounting medium supplemented with 1 ug/ml 4,6-diamidino-2-phenylindole (DAPI; Vector Laboratories, Inc.). Slides were examined on a Zeiss Axiovert 200 microscope with an AxioCam camera and ZEN Pro software (Carl Zeiss, Germany). For co-localization cells were analysed by confocal microscopy with a Leica TCS SP8 confocal laser scanning microscope and the Leica Application Suite X (LASX) software (Leica, Germany). Images were acquired as *z*-stacks (0.25 μm). Digital images were processed using the Omero Open microscopy environment (University of Dundee; https://www.openmicroscopy.org/omero/). In all cases, images for a specific analysis were acquired with identical settings.

### Protein turnover

To determine protein half-life, translation was blocked by addition of cycloheximide (100 μg/ml) and cells were harvested at various times by centrifugation (800 g for 10 min at 4°C). Cells were washed with ice-cold PBS, then resuspended in 1x SDS sample buffer (Thermo) and incubated at 70°C for 10 min. Samples were subjected to standard SDS-PAGE electrophoresis.

### Western blotting

Proteins were separated by electrophoresis on a 4-12% precast acrylamide Bis-Tris gel (Thermo) and transferred to PVDF membranes using the iBlot2 system (23 V, 6 min; Thermo). Non-specific binding was blocked using 5% (w/v) bovine serum albumin (BSA; Sigma) in Tris-buffered saline (pH 7.4) with 0.2% (v/v) Tween-20 (TBST). Membranes were incubated with primary antibodies diluted in TBST supplemented with 1% BSA overnight at 4°C. Antibodies were used at the following dilutions: rat anti-HA epitope IgG_1_ (clone 3F10; Sigma) at 1:5,000, rabbit anti-ISG75 (in house) at 1: 10,000, anti-mouse b-tubulin (clone KMX-1; Millipore) at 1: 10,000. Following five washes with TBST of 10 min each, membranes were incubated with secondary antibodies diluted in TBST supplemented with 1% BSA. Dilutions of horseradish peroxidase (HRP)-coupled secondary antibodies (Sigma) were anti-rat-HRP at 1: 10,000, anti-rabbit-HRP at 1: 10,000, anti-mouse-HRP at 1: 10,000. Detection was carried out by incubating membranes with ECL Prime Western Blotting System (Sigma) and GE healthcare Amersham Hyperfilm ECL (GE). Densitometry quantification was conducted using ImageJ software (NIH). For quantification using the Li-COR system (Li-Cor Bioscience, Lincoln NE), the following antibodies were diluted in Odyssey blocking buffer (Li-COR): goat anti-rabbit IgG: IR Dye680RD and goat anti-mouse or anti-rat IgG: IRDye800CW (Li-COR). All washes were with PBS supplemented with 0.5% Tween20. Quantitative Fluorescence signals were quantified on an Odyssey CLx Imager and processed using Li-COR software (Li-COR).

### Blue native PAGE (BN-PAGE)

BN-PAGE was performed using the NativePAGE Bis-Tris gel system (Thermo). Briefly, cells were washed three times with 1X PBS supplemented with protease inhibitor cocktail without EDTA (Roche) and solubilized in Native PAGE sample buffer supplemented with 10% glycerol, 1% n-dodecyl-b-d-maltoside, 1x protease inhibitor cocktail without EDTA (Roche), 100 μg/ml microccocal nuclease (NEB), and 1x microccocal nuclease buffer (NEB). Samples were incubated in solubilization buffer on ice for 30 min and centrifuged (13,000 g at 4 °C, 30 min). The resulting supernatants were fractionated on precast 4-16% BN gradient gels (Thermo).

### Affinity isolation

Ubiquitylated proteins were isolated using the UbiQapture-Q kit (Enzo Life Sciences, Farmingdale, New York, USA) according to the manufacturer’s instructions. Ubiquitylated proteins were isolated from a total of 1 × 10^7^ cells lysed with TEN buffer (150 mM NaCl, 50 mM Tris-HCl, pH 7.4, 5 mM EDTA, 1% Triton X-100), supplemented with 100 mM N-ethylmaleimide (NEM) to inhibit deubiquitinase activity [36]. A total of 40 μl of UbiQapture-Q^TM^ matrix was pre-equilibrated in TEN buffer and incubated with cell lysates (200 μl) by rotating at 4°C overnight. After washing five times, captured proteins were eluted with 2x SDS-PAGE sample buffer containing 10mM DTT. Samples were resolved in 4-12% acrylamide gels, transferred onto PVDF membranes and analyzed by Western blotting using anti-HA antibody in blocking buffer (TBST supplemented with 1% BSA).

### Molecular modelling

A homology model of TbAQP2 tetramer (residues 68-312) was built using Modeller (version 9.20) [37,38] using as template the crystal structure of the *Homo sapiens* AQP10 (PDB code 6f7H) [39]. The N-terminus (residues 1-59) could not be modelled due to predicted flexibility and low sequence similarity. *T. brucei* AQP2 has 33% identity compared to *Homo sapiens* AQP10. Multiple sequence alignments were performed using T-Coffee [40] and ClustalW [41]. The geometries of the homology model were refined using Maestro and verified using PROCHECK [42] The resulting Ramachandran plots indicated a good model quality with 93% of the residues in most favoured regions. A second model displaying K147R and K234R mutations in each monomer was generated following the same protocol. Both models were refined using all-atom molecular dynamics (MD) simulations with Desmond [43]. Each system was embedded as a tetramer in a periodic POPC lipid bilayer generated with “System Builder” in Maestro and solvated in aqueous 150mM KCl. The OPLS3e force field was used to further improve the resulting molecular model [44]. The cut-off distance for non-bonded interactions was 9 Å. The SHAKE algorithm was applied to all bonds involving hydrogens, and an integration step of 2.0 fs was used throughout [45]. The systems were simulated with no restraints at constant temperature (300K) and pressure (1atm) for 100ns. Protein structures and MD trajectories were visually inspected and analysed using the molecular visualization programs PyMOL, VMD [45] and Maestro [43].

## Results

### TbAQP2 forms stable tetrameric complexes in the bloodstream form of T. brucei

TbAQP2 (Tb927.10.14170) is critical for water and glycerol transport activity as well as sensitivity to diamidines and melaminophenyl arsenicals in African trypanosomes [13,23,33]. A recent study suggested that pentamidine may be a nanomolar ligand, rather than a transport substrate of TbAQP2 [22] and that endocytosis of TbAQP2 might be important for pentamidine transport. However, the intracellular and surface trafficking pathways of AQPs in trypanosomes have not been elucidated. Central to endocytosis and protein sorting of many surface membrane proteins is ubiquitylation [46], and ubiquitylation of the type I surface-localised invariant surface glycoproteins 65 and 75 (ISG65 and ISG75) is essential for internalization and degradation in the lysosome [33,47,48].

To determine whether ubiquitylation is involved in trafficking and turnover of polytopic surface proteins in trypanosomes we addressed whether TbAQP2 is ubiquitylated *in vivo*. We generated *T. brucei* cell lines expressing TbAQP2 tagged at the N- (^3×HA^AQP2) or C-terminus (AQP2^3×HA^) (**Fig 1A and B**) using an *aqp-*null cell line [19] as chassis to prevent heterologous interaction with endogenous AQPs. The hemagglutinin (HA) tag was selected as it lacks lysine residues (as opposed to more bulky tags such as GFP) and therefore is incapable of becoming ubiquitylated and interfering with data interpretation. Both constructs co-localized with ISG75 at the posterior end of the cell, consistent with the location of native AQP2 at the flagellar pocket (**Fig 2A**). ^3×HA^AQP2 is predominantly detected as two forms by immunoblotting after SDS-PAGE: a ∼38 kDa form, consistent with the monomeric form, and a >120 kDa form, likely a homotetramer (**Fig 2B, lower panel**), as previously reported for other AQPs [16,49–51]. In sharp contrast, AQP2^3×HA^ was found as two main species of ∼35 kDa and ∼38 kDa, with no tetrameric form detected (**Fig 2B, lower panel**). However, under native conditions, both constructs are organized as high molecular weight complexes of ∼480 kDa, consistent with a 4×4 conformation under native conditions (**Fig 2B, upper panel**).

**Figure 2.**
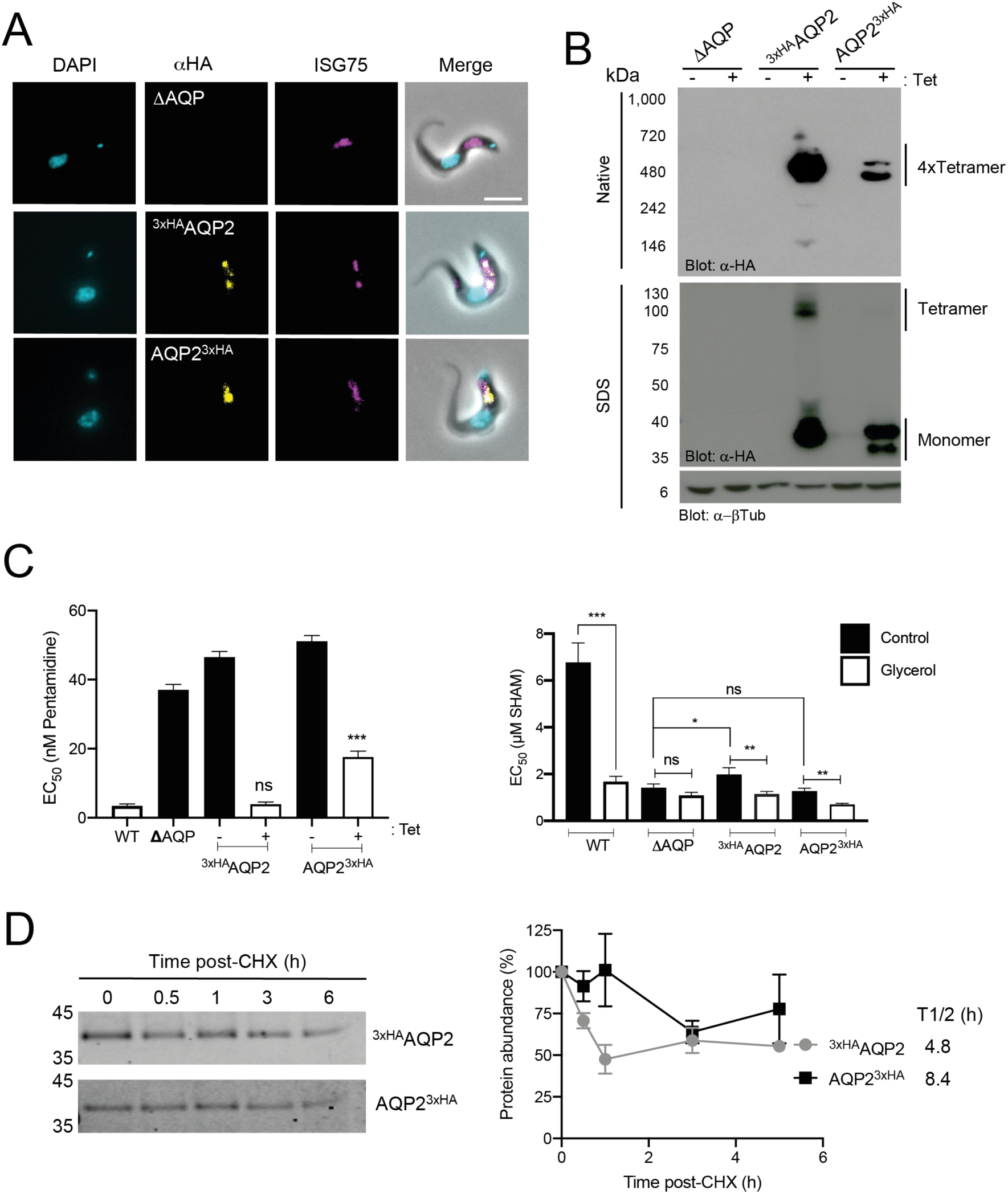
Characterisation of tagged TbAQP2. **A)** Fluorescence microscopy of *T. brucei* 2T1 cells expressing tetracycline-regulated N- or C-terminal tagged AQP2 (^3xHA^AQP2 or AQP2^3xHA^, respectively, in yellow). These proteins localise similar to ISG75 (magenta) at the flagellar pocket/endosomes. The triple *aqp-*null *T. brucei* 2T1 cells (*Δ*AQP) were also included as control. Scale bar 5 μm. **B)** Tet-regulated expression of N- or C-terminal HA-tagged AQP2. Both native-PAGE (upper panel) and SDS-PAGE (lower panel) αHA blots are shown. α−β tubulin was used as loading control. The presence of the different oligomeric species is indicated in the right-hand side of the panel. Note the presence of a high molecular weight form under SDS-PAGE in ^3xHA^AQP2 but not AQP2^3xHA^. The triple *aqp-*null *T. brucei* 2T1 cells (*Δ*AQP) were also included as control. **C)** EC_50_ values for pentamidine **(left panel)** or salicylhydroxamic acid (SHAM; **right panel**) with or without 5 mM glycerol following expression of either ^3xHA^AQP2 or AQP2^3xHA^. For multiparametric ANOVA, we compared the average values (n = 4 independent replicates) from wild type *T. brucei* 2T1 cells as reference for pentamidine, or from *aqp-*null cells for SHAM. * *p*<0.01, ** *p*<0.001, *** *p*<0.0001 from four independent replicates. **D) Left panel;** Representative western blotting (n = 3 independent replicates) from protein turnover assay monitored by cycloheximide (CHX) treatment in *T. brucei* 2T1 cells expressing either ^3xHA^AQP2 (upper panel) or AQP2^3xHA^ (lower panel). **Right panel**; Protein quantification from western blotting analysis in left panel for either ^3xHA^AQP2 (black square) or AQP2^3xHA^ (grey circles). Results are the mean ± standard deviation of three independent experiments (n = 3 independent replicates). The estimated half-life (t_1/2_) was calculated based on regression analysis using PRISM.

Thus, whereas ^3×HA^AQP2 is readily detectable as a stable tetramer, even under harsh conditions, AQP2^3×HA^ is comparatively less stable in its tetrameric form (**Fig 2B**), likely indicating interference by the C-terminal HA tag to oligomerization and/or tetramer stability. To determine the glycerol transport capacity of these proteins, we inhibited the activity of the trypanosome alternative oxidase (TAO) with salicylhydroxamic acid (SHAM) [21]. Inhibition of TAO leads to increased intracellular glycerol, building up to toxic levels that can only be prevented by export *via* a glycerol transporter such as AQP. Therefore, the absence of functional AQPs renders cells highly susceptible to SHAM. Consistent with stability of the AQP2^3xHA^ oligomeric form, expression of the ^3×HA^AQP2 construct in the *aqp-null* background restored sensitivity to pentamidine and glycerol transport comparable to wild type cells, whereas AQP2^3×HA^ only partly rescued these phenotypes (**Fig 2C and S1 Fig**). Both ^3×HA^AQP2 and AQP2^3×HA^ have long half-lives (>4h) (**Fig 2D**) indicating that impaired transport activity of AQP2^3xHA^ is unlikely due to altered turnover or structure. Therefore, although introduction of HA epitopes to either terminus does not alter localization, only the N-terminal tagged form assembles stable oligomeric structures and fully functional TbAQP2. Therefore, we selected to focus on ^3xHA^AQP2.

### TbAQP2 is ubiquitylated and degraded by the lysosome

Next, we sought evidence that TbAQP2 is ubiquitylated. Cycles of protein ubiquitylation and deubiquitylation are important for controlling the cell surface composition of trypanosomes [25,33,52]. Both proteasome-dependent and lysosome-mediated protein degradation are generally initiated by covalent attachment of one or more ubiquitin moieties to a substrate protein [53], and we rationalized that inhibiting these degradative systems would increase the overall abundance of ubiquitylated intermediates.

We observed high molecular weight adducts when cells expressing ^3×HA^AQP2 were treated with either ammonium chloride (lysosomal activity inhibitor) or MG132 (proteasome inhibitor) (**Fig 3A**), likely representing ubiquitylated intermediates *en route* to degradation. Subsequent western blotting identified a predominant band of ∼55 kDa reactive to anti-ubiquitin antibody upon immunoprecipitation with anti-HA magnetic beads, consistent with the addition of ubiquitin to TbAQP2 (∼38 kDa for unmodified protein (**Fig 3B**). To corroborate these results, we performed an affinity isolation using a commercial ubiquitin binding domain (UBD) resin followed by western blotting with anti-HA antibody. This revealed unmodified monomer together with high molecular weight adducts, likely representing TbAQP2 with various numbers of ubiquitin conjugates; the latter clearly represents a small fraction of total AQP2 expressed in these cells (**Fig 3C**). Interestingly, we noted a band of around ∼40 kDa, likely corresponding to monoubiquitylated TbAQP2 (**Fig 3C**). Collectively, these results indicate that TbAQP2 is modified by ubiquitin in the bloodstream form of *T. brucei*.

**Figure 3.**
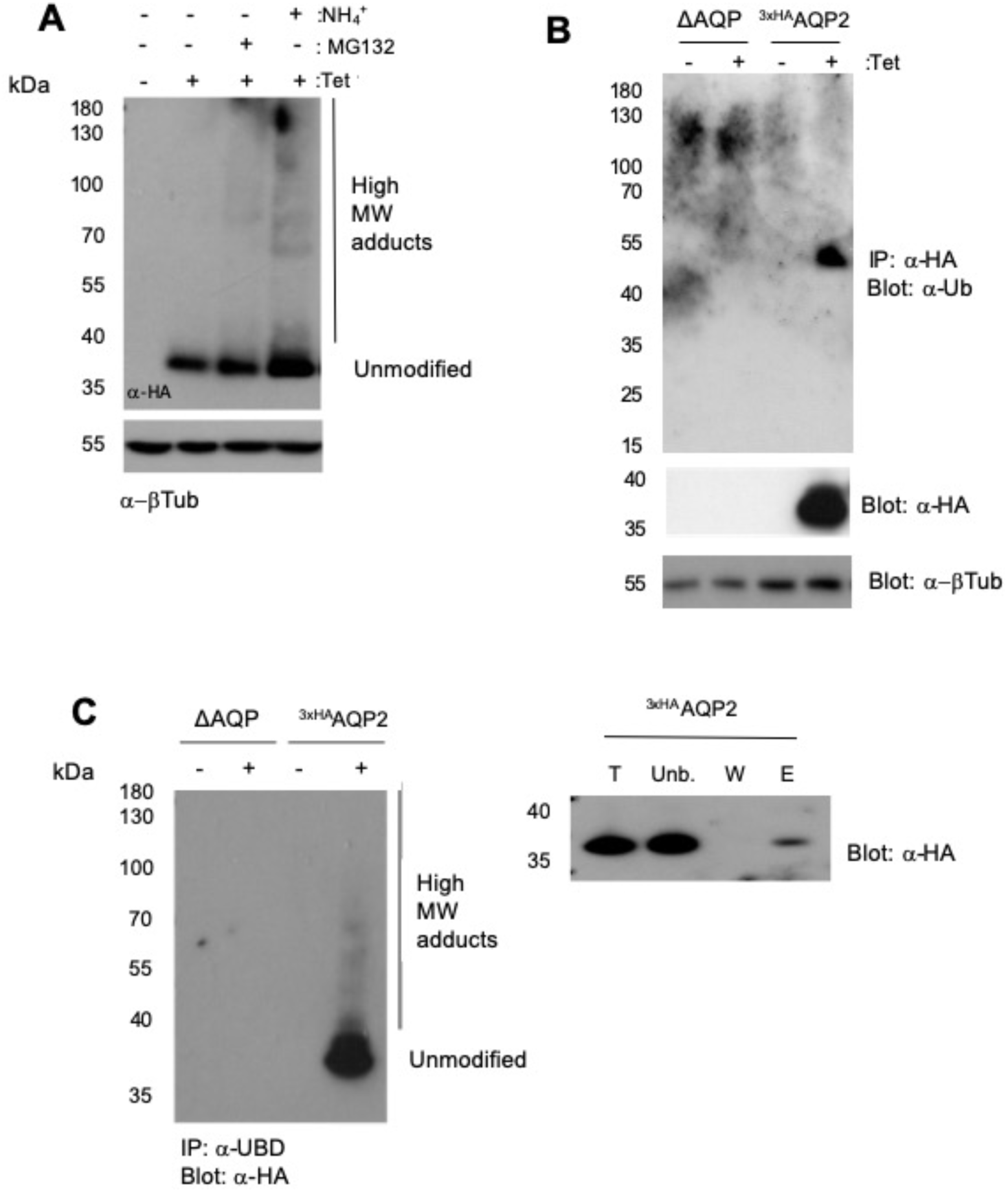
TbAQP2 is ubiquitylated in *T. brucei*. **A)** Cells expressing ^3xHA^AQP2 were treated with either NH_4_^+^Cl (10 mM) or MG132 (25 μM) for 1h prior to harvesting. Cell lysates were resolved in a 4-12% acrylamide gel and detected with anti-HA antibody by western blotting. The intensity of anti-β tubulin was used as loading control. **B)** Immunoprecipitation of *Δaqp* or ^3xHA^AQP2 cell lysates with anti-HA beads followed by anti-ubiquitin detection by western blotting. An anti-HA blot was also included to confirm protein expression upon induction with tetracycline. Anti-β tubulin was used as loading control. **C)** As in (B), but immunoprecipitation conducted using ubiquitin capture matrix and analysed by western blotting (left panel). The total (“T”), unbound (“Unb.”), wash (“W”), and elution (“E”) fractions were resolved by SDS-PAGE electrophoresis and analysed with anti-HA immunoblotting (right panel).

Next, we sought to investigate the mechanisms by which TbAQP2 is degraded. Imaging suggested that TbAQP2 is predominantly located at the flagellar pocket together with ISG75, but a proportion is also in close proximity to early endosomes (positive for Rab5A and Rab5B) but less so for recycling endosomes (Rab11) (**Fig 4A**) suggesting transit of TbAQP2 through early endosomes. Moreover, TbAQP2 displayed strong overlap with p67, a lysosomal marker, suggesting that TbAQP2 is delivered to the lysosome *via* endocytosis (**Fig 4A**). Similar observations were made with cells expressing AQP2^3×HA^ (**S2 fig**), once more indicating that the C-terminal tag does not impair trafficking but rather hinders oligomerisation. Further, pulse-chase analysis showed that ISG75 has a half-life of ∼3.6 h, consistent with earlier studies [33,47,48], whereas TbAQP2 displays bimodal behaviour with approximately half rapidly turned over in <1 h, with the remaining fraction is more stable, with a half-life of ∼6 h (**Fig 4B**), which together with partial juxtaposition with Rab11, suggests possible recycling. To determine whether TbAQP2 is degraded in the lysosome or the proteasome we treated cells with bafilomycin A1 (BafA1; inhibitor of the lysosomal v-ATPase) or MG132 (canonical proteasome inhibitor with broad-range inhibitory capacity against serine proteases and calpain-like proteases [54]). In untreated cells, TbAQP2 was reduced by ∼50% after 1 h as expected, but in cells treated with BafA1 or MG132, less that 20% of the protein was degraded (**Fig 4C**). It is important to note that MG132 can also impair degradation of proteins delivered to the lysosome as it acts as a broad range inhibitor for lysosome-specific proteases [54]. Overall, these data indicate that TbAQP2 is ubiquitylated and delivered to the lysosome for degradation, *albeit* with a pool of longer-lived protein that may constitute a recycling population.

**Figure 4.**
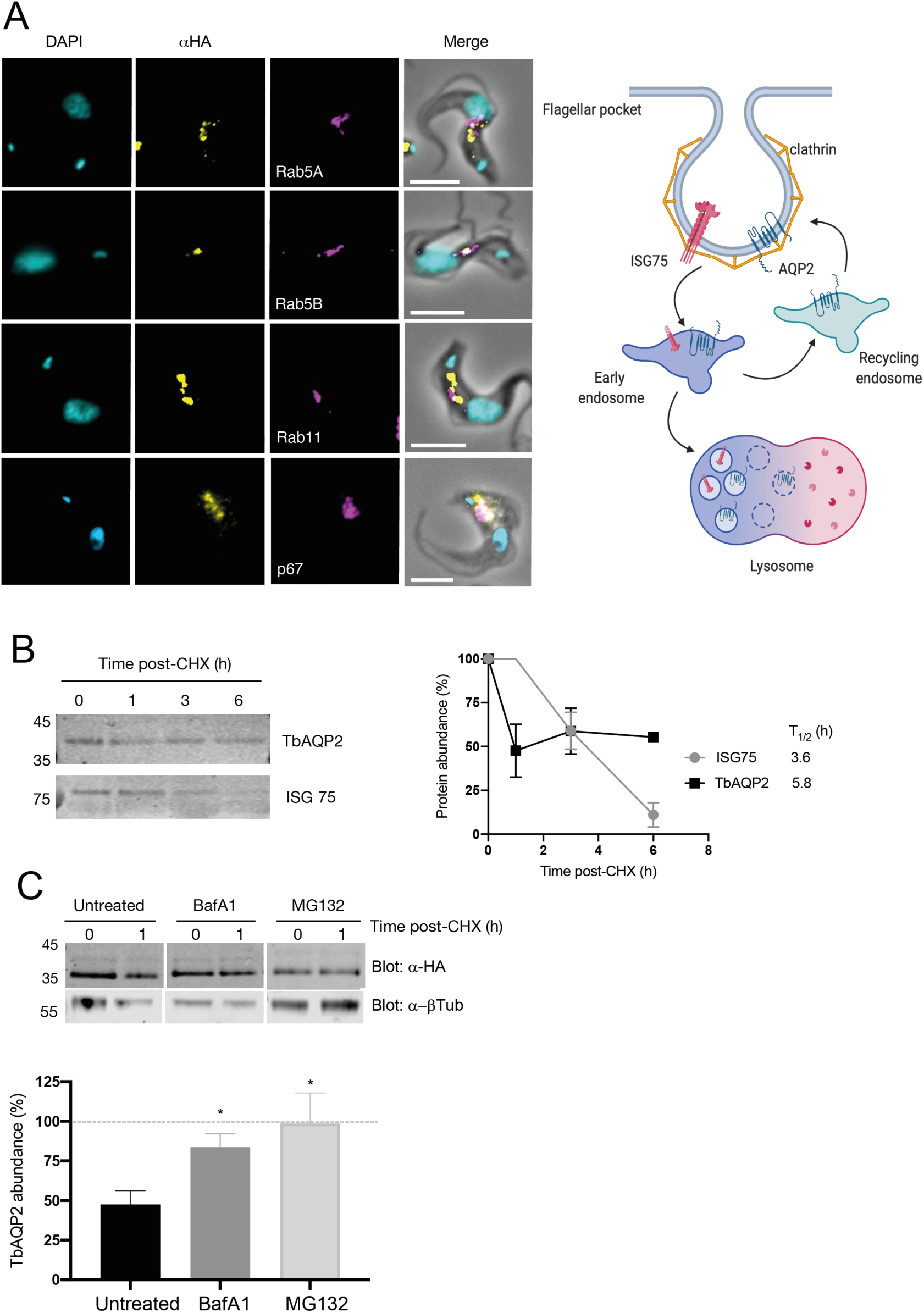
TbAQP2 transits through the endosomal compartment and is efficiently delivered and degraded in the lysosome. **A)** Cell lines expressing a tetracycline-regulated copy of ^3xHA^AQP2 (Alexa Fluor 488; yellow) were co-stained with anti-TbRab5a and anti-TbRab5b (early endosomes), anti-TbRab11 (recycling endosomes), and anti-p67 (lysosome). All endosomal and lysosomal markers were labelled with secondary antibodies coupled to Alexa Fluor 568 (magenta). DAPI (cyan) was used to label the nucleus and kinetoplast. Scale bars 5 μm. A schematic depiction of the results from confocal microscopy is included in the right panel, generated with BioRender. **B) Left panel**; Protein turnover was monitored by cycloheximide (CHX) treatment. Cells were harvested at various times and the protein level monitored by immunoblotting. ISG75 was included as a control. **Right panel**; Quantification for ISG75 and ^3xHA^AQP2. Results are the mean ± standard deviation of three independent experiments. **C) Upper panel**; As in (B), but cells were untreated or exposed to 100 nM of bafilomycin A1 (BafA1), or to 25 μM of MG132 for 1 h prior to harvesting. Cell lysates were resolved by SDS-PAGE followed by western immunoblotting using anti-HA antibody. **Lower panel;** Quantification from three independent experiments - dotted line represents 100% (signal at 0h). Data presented as mean ± standard deviation (n= 3 independent replicates). Statistical analysis was conducted using *t* test; * *p*<0.01 and the signal from untreated cells at 1 h as reference.

### Intracellular N-terminal lysine residues are essential for oligomerisation and channel function of TbAQP2

Predictions of TbAQP2 topology [55] suggested cytosolic localisations for both N- and C-termini, as is known for the mammalian orthologues (**S3 Fig,** [15,56]). AQP2 has five lysine residues that are exposed to the cytosol, at positions 19, 45, 54, 147, and 234 (**Fig 1B**). To better understand the potential ubiquitylation sites in TbAQP2, we used UbPred (http://www.ubpred.org) [57] to predict lysine residues as candidate ubiquitin acceptors. UbPred suggested that lysine residues in position 19, 45, and 54 are potential ubiquitylation sites in TbAQP2, with prediction scores of 0.65, 0.73, and 0.88, respectively. All three residues are located within the N-terminal cytoplasmic region of AQP2 (**Fig 1B**).

To dissect the contribution of these residues to TbAQP2 localisation and function, we generated a cell line expressing N-terminally tagged AQP2 in which all three of these lysine residues were simultaneously mutated (AQP2^3K>R^). Unexpectedly, while the wild-type protein located in the posterior end of the cell, AQP2^3K>R^ was mislocalized (**Fig 5A**) and failed to restore pentamidine sensitivity and glycerol transport (**Fig 5B**). Furthermore, whereas AQP2^WT^ co-localises with ISG75 at the posterior end of the cells, AQP2^3K>R^ was retained in the endoplasmic reticulum (ER), as suggested by co-localisation with the ER marker TbBiP (**Fig 5C**). Blue native-PAGE indicated that whereas AQP2^WT^ forms two high molecular weight complexes (∼480 kDa and ∼120 kDa), AQP2^3K>R^ did not oligomerise (**Fig 5D and S4 Fig**). Moreover, AQP2^3K>R^ is highly unstable and turned over faster than AQP2^WT^ and in an MG-132 selective manner (**Fig 5E**). We conclude that K19, K45 and K54 are essential for TbAQP2 folding and hence anterograde trafficking and that their replacement by arginine triggers entry into an ER-associated degradative (ERAD) pathway [58–60].

**Figure 5.**
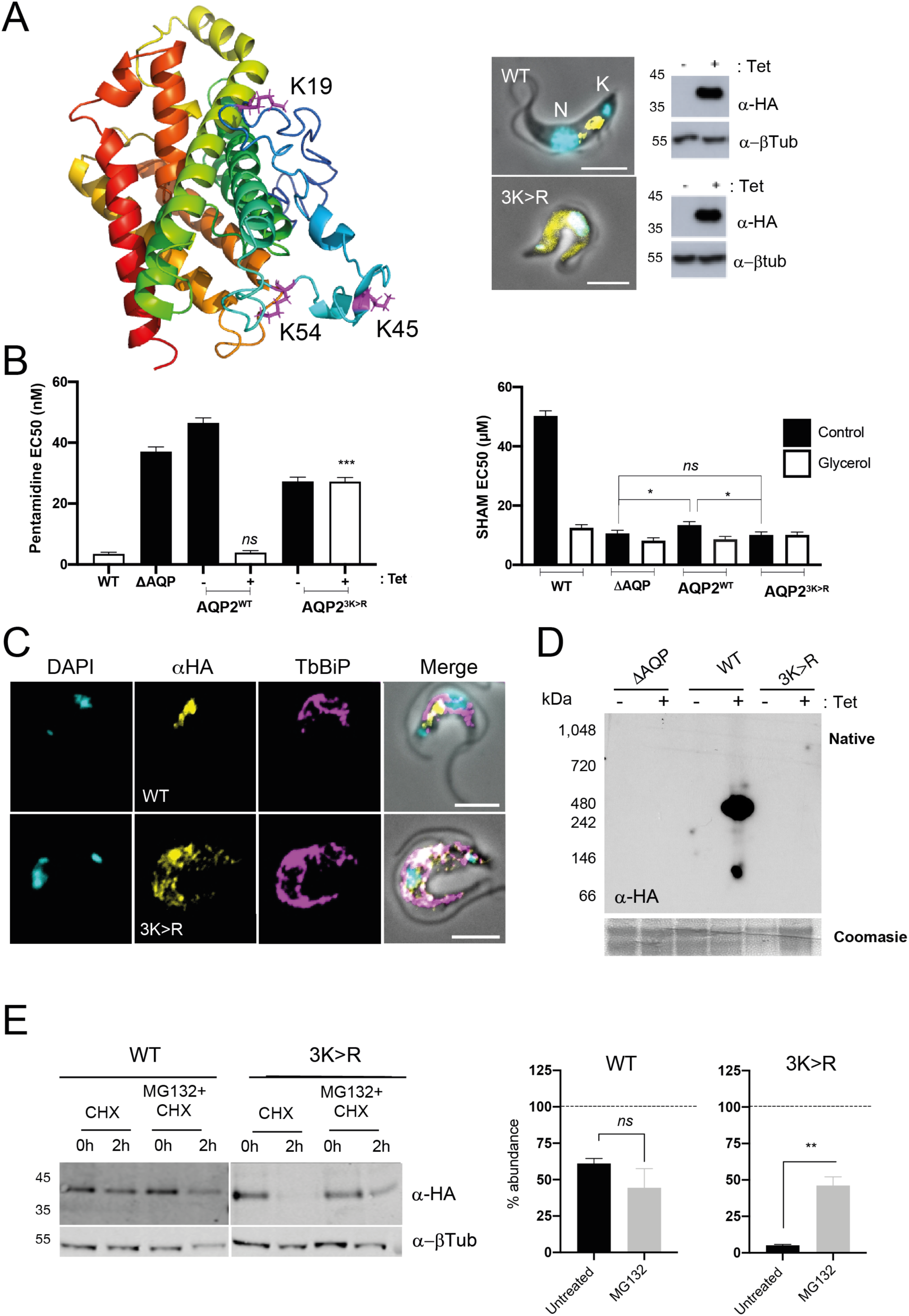
N-terminal lysine residues in the N-terminal cytoplasmic tail are important for protein stability, oligomerisation, and anterograde transport. **A) Left panel**; Structural predictions of ^3xHA^AQP2 generated with i-Tasser, indicating the three N-terminal lysine residues (magenta) mutated in AQP2^3K>R^. The 3xHA tag has been omitted for simplicity. **Right panel**; Fluorescence microscopy of cells expressing N-terminal HA-tagged wild type AQP2 (AQP2^WT^) or lysine mutant AQP2^3K>R^. Both proteins are shown in yellow. DAPI (cyan) was used to label the nucleus (N) and the kinetoplast (K). Scale bars, 5 μm. Western blot of cell lysates upon induction with tetracycline are also included. **B)** EC_50_ values for pentamidine (**left panel**) and salicylhydroxamic acid (SHAM) with or without 5 mM glycerol (**right panel**) following recombinant expression of either AQP2^WT^ or AQP2^3K>R^ with a tetracycline-regulated (Tet-on) copy in *T. brucei* 2T1 bloodstream forms. Multiparametric ANOVA calculated as for Figure 4 (*N* = 3 independent replicates). **C)** Cell lines expressing AQP2^WT^ or AQP2^3K>R^ (Alexa Fluor 488; yellow) were co-stained with anti-BiP (endoplasmic reticulum marker). All markers were labelled with secondary antibodies coupled to Alexa Fluor 568 (magenta). DAPI (cyan) was used to label the nucleus and the kinetoplast, as indicated in (A). Scale bars, 5 μm. **D)** Native-PAGE immunoblot of total cell lysates expressing either AQP2^WT^ or AQP2^3K>R^. Coomassie blue staining of the same fractions was used as loading control. The triple *aqp-*null *T. brucei* 2T1 cells (*Δ*AQP) were also included as control. **E) Left panel;** Protein turnover monitored as in Figure 4 for AQP2^WT^ or AQP2^3K>R^. Cells were either untreated or treated with 25 μM MG132 for 1 h prior to harvest. Cells were harvested at 0 hours and 2 h post-CHX treatment, and lysates analysed by immunoblotting. α−β tubulin was used as loading control. **Right panel**; Protein quantification representing the mean ± standard deviation (n = 3 independent replicates). Dotted line represents 100% (signal in untreated samples). Statistical analysis was conducted using the signal from untreated cells at 2 h as reference group. ** *p*<0.001, ns = not significant, using a *t* test.

### Site-directed mutation of cytoplasmic lysine residues of TbAQP2 leads to protein instability

To determine whether the effects observed for AQP2^3K>R^ could be attributed to a single lysine residue we generated a construct in which all cytoplasmic lysine residues were mutagenized to arginine (AQP2^5K>R^) (**Fig 1B**). Using this construct as template, we reverted each lysine individually using site-directed mutagenesis, generating cell lines expressing N-terminally tagged mutant TbAQP2 with only one lysine residue reinstated (AQP2^R19K^, AQP2^R45K^, and AQP2^R54K^). None of these mutants formed oligomers (**S4A Fig.**) and were retained in the ER, as demonstrated by co-localisation with TbBiP (**Fig 6A**). Moreover, AQP2^5K>R^, AQP2^R19K^, AQP2^R45K^, and AQP2^R54K^ turn over faster than AQP2^WT^ and are stabilised by MG-132 (**Fig 6B and Table 1**), consistent with the absence of detection by BN-PAGE analysis and the lack of sensitivity to pentamidine and glycerol transport observed in these mutants (**Fig 6C**). Similar results were obtained with AQP2^R234K^ (**S4F Fig**). K234 is located within the TM4-TMD5 loop, an important feature of TbAPQ2 as this loop is predicted to interact with the TM4-TM5 loop of the neighbouring subunit to create a large oligomerization interface (**S4G Fig, left panel**).

**Figure 6.**
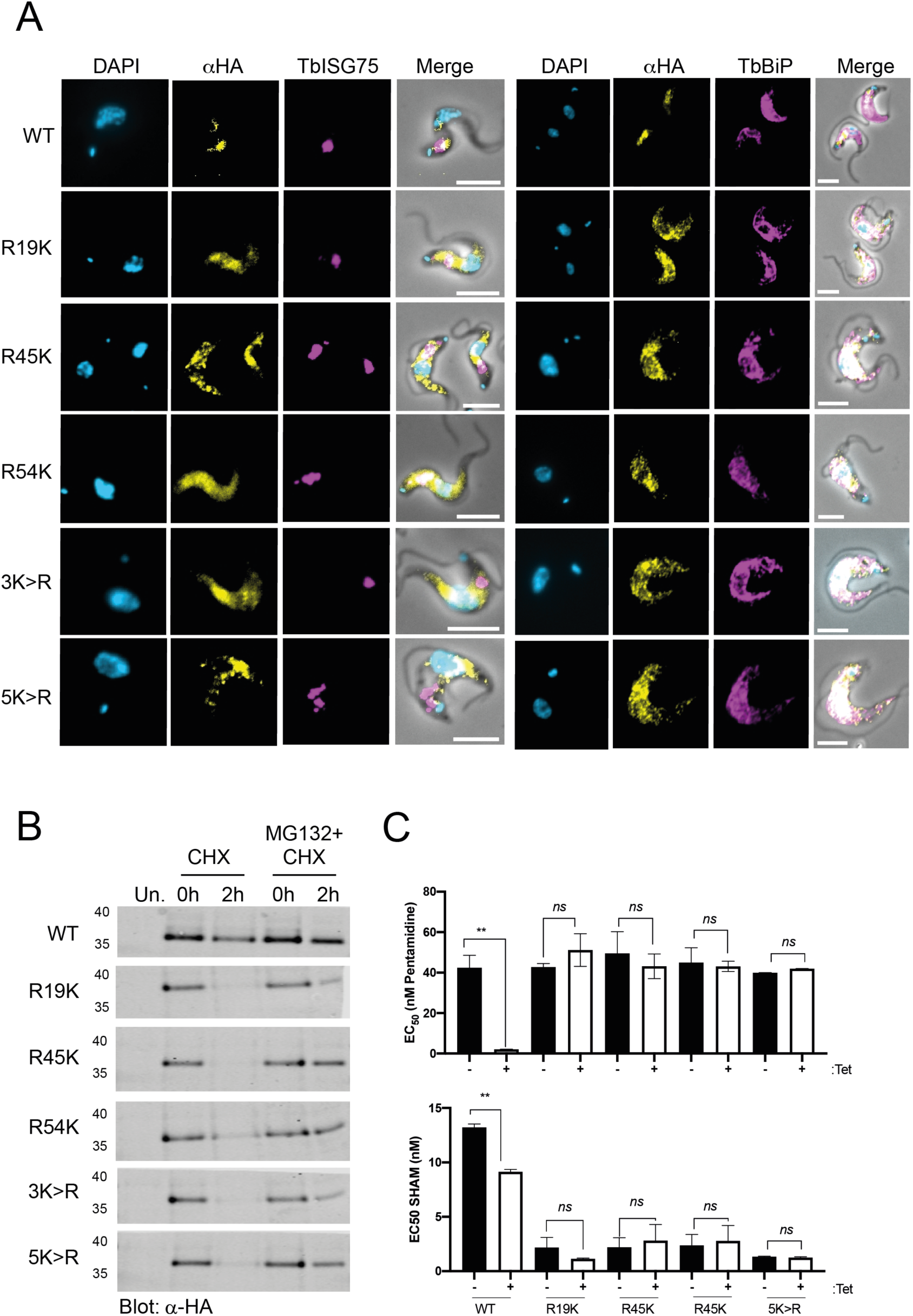
Requirement for cytoplasmic-oriented lysine residues for AQP2 stability and trafficking. **A)** Cell lines expressing a tetracycline-regulated copy of the constructs mentioned in (A) (Alexa Fluor 488; yellow) were co-stained with either αBiP (ER) or αISG75 (localises to flagellar pocket/endosome), both stained with secondary antibodies coupled to Alexa Fluor 568 (magenta). DAPI (cyan) was used to label the nucleus and the kinetoplast. Scale bars, 5 μm. **B)** Representative western blot (n = 3 independent replicates) of protein turnover monitored by cycloheximide (CHX) treatment followed by pulse-chase of cells expressing the constructs in (A). Cells were either untreated or treated with 25 μM MG132 for 1 h prior to harvest. Cells were harvested at 0 hours and 2 h post-CHX treatment and analysed by immunoblotting. Uninduced controls (“Un.”) were also included. **C**) EC_50_ values (average ± standard deviation; n = 3 independent replicates) of pentamidine (upper panel) and salicylhydroxamic acid (SHAM) with or without 5 mM glycerol (lower panel) following recombinant expression of either AQP2^WT^, AQP2^5K>R^, or single arginine-to-lysine AQP2 mutants (AQP2^R19K^, AQP2^R45K^, and AQP2^R54K^). Statistical test for significance was conducted using a pairwise *t* test comparison with uninduced cell lines. * *p*<0.01, ** *p*<0.001, *** *p*<0.0001.

**Table 1.** Summary of the impact of cytoplasmic lysine mutagenesis on TbAQP2 localisation and function.

| Protein | Localisation | Protein abundance post-CHX<sup>2</sup> |  | Proposed degradation pathway | EC<sub>50</sub> pentamidine (nM)<sup>3</sup> | EC<sub>50</sub> SHAM (μM)<sup>3</sup> | EC<sub>50</sub> SHAM + 5 mM glycerol (μM)<sup>3</sup> |
| --- | --- | --- | --- | --- | --- | --- | --- |
|  |  | Untreated | + MG132 |  |  |  |  |
| **AQP2<sup>WT</sup>** | Flagellar pocket | 61.05 ± 3.43% | 44.42 ± 13.15% | Lysosome | 3.29 | 1.96 | 1.16 |
| **<sup>1</sup>AQP2<sup>R19K</sup>** | Endoplasmic reticulum | 14.42 ± 9.5% | 16.11 ± 7.1% | ERAD<sup>1</sup> | 51.18 | 1.92 | 1.12 |
| **AQP2<sup>R45K</sup>** | Endoplasmic reticulum | 1.3 ± 0.93% | 34.3 ± 4.65% | ERAD | 43.16 | 1.99 | 2.36 |
| **AQP2<sup>R54K</sup>** | Endoplasmic reticulum | 10.84 ± 0.5% | 41.05 ± 12.5% | ERAD | 43.10 | 2.10 | 2.38 |
| **AQP2<sup>R234K</sup>** | Endoplasmic reticulum | 7.67 ± 1.9% | 24.91 ± 5.12% | ERAD | 39.95 | 1.28 | 2.34 |
| **AQP2<sup>3K>R</sup>** | Endoplasmic reticulum | 5.16 ± 0.62% | 46.18 ± 5.95% | ERAD | 27.20 | 10.10 | 10.14 |
| **AQP2<sup>5K>R</sup>** | Endoplasmic reticulum | 16.8 ± 6.2% | 37.5 ± 9.5% | ERAD | 41.98 | 1.34 | 1.25 |

<sup>1</sup>AQP2<sup>R19K</sup> did not show a significant accumulation upon MG132 treatment, but co-localized with TbBiP, indicating probable ERAD-mediated turnover. <sup>2</sup>Protein abundance was calculated 2h post-treatment with cycloheximide (CHX) and expressed as percent of protein abundance compared to protein signal before treatment ("time 0h"). <sup>3</sup>Estimated EC<sub>50</sub> values from cells induced with tetracycline, 1 μg/ml for 24h.

Molecular dynamics (MD) simulations of Tb^WT^ and Tb^K147R/K234R^ suggest that the K234R mutation will likely have a notable effect on the position of the TM4-TM5 loop, hampering oligomerization (**S4G Fig**). It is important to note that we failed to successfully express AQP2^R147K^ despite multiple independent transfections. Residue 147 is located between TMD4 and TMD5, potentially indicating that mutation of this residue leads to a far more unstable protein than the other constructs, and in good agreement with the MD simulations. In TbAQP2^WT^, K147 is predicted to interact with Y151 and N70 on TMD1 and maintain the TMD3-TMD1 interface (**S4G Fig, right panel).** On the other hand, simulations of TbAQP2^K147R/K234R^ showed a significant conformational change of TMD1 and TMD3 as a result of establishment of a salt bridge between R234 and D73 on TMD1. This conformational change on TMD1 would impact both the conformation of the N-terminal tail and the dimerization interface with TMD6 from the neighbouring subunit, providing a rationale for the instability observed in this mutant. Consistent with a lack of oligomerization and rapid turnover all of these constructs failed to restore pentamidine sensitivity or glycerol transport (**Fig 6C and Table 1**).

### Chimerization of TbAQP2 impairs stability, localization and function

*T. brucei* possesses three AQP paralogues [21,23]. Of these, TbAQP2 and TbAQP3 are tandem open reading frames located on chromosome 10 and share >70% protein identity [13]. Chimerisation of the loci encoding TbAQP2 and TbAQP3 causes resistance to pentamidine and melarsoprol [27,29,32,56]. Interestingly, although in some cases the selectivity pore is mutated, many chimeric AQP2/3 alleles do not have altered amino acids in the selectivity pore, but rather replacement of TMD regions with sequences from TbAQP3 (**Fig 1C**), [29–32]. Moreover, the AQP2/3 chimeras characterised so far display a subcellular localisation resembling TbAQP3 at the plasma membrane, in contrast with an expected flagellar pocket localisation for TbAQP2 (**Fig 2A**) [27]. However, it is unclear if TbAQP2 chimerisation impacts additional features beyond subcellular localisation.

We generated cell lines expressing tetracycline-regulated N-terminal tagged TbAQP1, TbAQP2, TbAQP3 and the chimeric AQP2/3 40AT (40AT) (**Fig 1C**), isolated from relapse patients from the Democratic Republic of Congo [29]. One of the main structural features of this chimera is replacement of the sixth *trans*-membrane and C-terminus of TbAQP2 with the corresponding sequence of TbAQP3 (**Fig 1C**) [29]. Additionally, to simulate other chimeric AQP2/3 proteins identified in laboratory strains and field isolates, we generated AQP2 mutants where TMD4 (AQP2^TMD4^) and TMD5 (AQP2^TMD5^) are individually replaced by the corresponding TMDs from TbAQP3 (**Fig 1C**). Apart from the AQP2^TMD5^ construct, none of these constructs alter the amino acid composition of the selectivity filter of TbAQP2 (**S5 and S6 Fig**). Whereas we readily expressed AQP2^TMD4^, we failed to obtain positive clones for AQP2^TMD5^, despite multiple attempts. We observed that TbAQP2 colocalised with ISG75, as expected, as well as TbAQP1 which seems to localise in close proximity to ISG75, whereas TbAQP3 localises mainly to the cell surface as previously reported [13,21] (**Fig 7A, left-hand panel**). Conversely, AQP2^TMD4^ and the clinical chimera 40AT displayed a distinct localisation in proximity with TbBiP (**Fig 8A**). Western blotting showed that under reducing conditions, all these constructs are readily detected as monomers of ∼35-38 kDa (**Fig 7A, right-hand panel**). Under native conditions, both TbAQP1 and TbAQP2 can be readily detected as *n*-dodecyl b-D-maltoside (DDM)-soluble forms of ∼480 kDa species, consistent with the proposed 4×4 organization, whereas we failed to observe such complexes for TbAQP3, 40AT, and AQP2^TMD4^ (**Fig 7B**). TbAQP3, AQP2^TMD4^ and 40AT are turned over more rapidly (<1 h) than TbAQP1 and TbAQP2 (**Fig 7D and Table 2**), explaining the lack of glycerol transport in cells expressing these constructs (**Fig 7C**).

**Figure 7.**
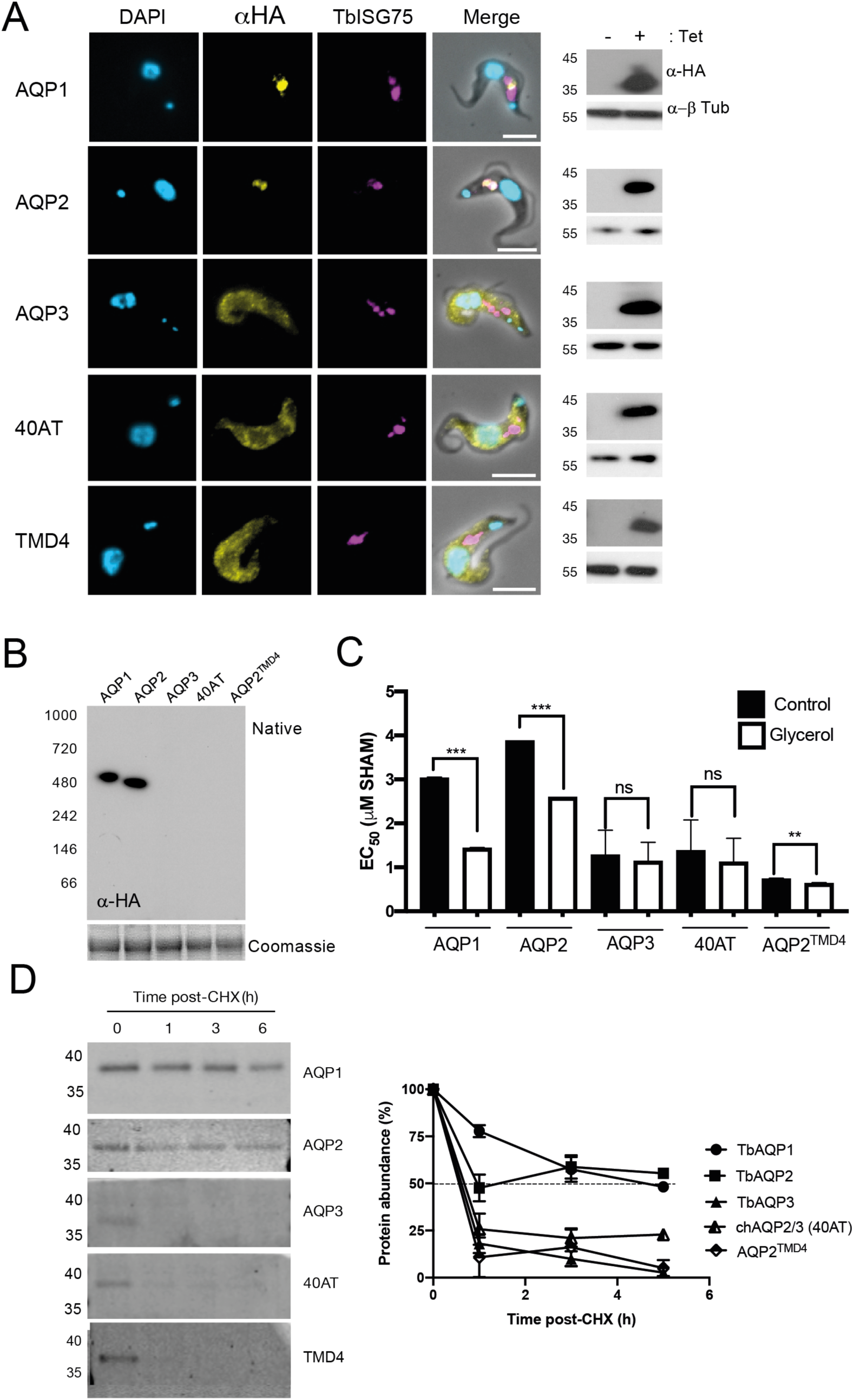
Chimerisation of TbAQP2 leads to mislocalisation, reduction in glycerol transport activity and rapid turnover. **A)** Cell lines expressing N-terminal HA-tagged TbAQP1, TbAQP2, TbAQP3, field-isolate chimeric AQP2/3 (40AT) or a single TMD mutant (AQP2^TMD4^) (Alexa Fluor 488; yellow) co-stained with anti-ISG75 (magenta). DAPI (cyan) was used to label the nucleus and the kinetoplast. Scale bars 5 μm. Western immunoblotting analysis from lysates of cells expressing these constructs are also included. Anti-β tubulin was used as loading control. **B)** BN-PAGE immunoblot of total cell lysates expressing the constructs in (A). Coomassie blue staining of the same fractions was used as loading control. **C)** EC_50_ values (average ± standard deviation; n = 3) for salicylhydroxamic acid (SHAM) with or without 5 mM glycerol following recombinant expression of the constructs in (A). **D) Left panel**; Representative western blotting (n = 3 independent replicates) analysis of protein turnover monitored by cycloheximide (CHX) treatment followed by pulse-chase of cells expressing the constructs in (A). **Right panel**; Protein quantification representing the mean ± standard deviation of three independent experiments. Dotted line represents 50% of protein abundance. Data presented as mean ± standard deviation (n = 3 independent replicates).

**Figure 8.**
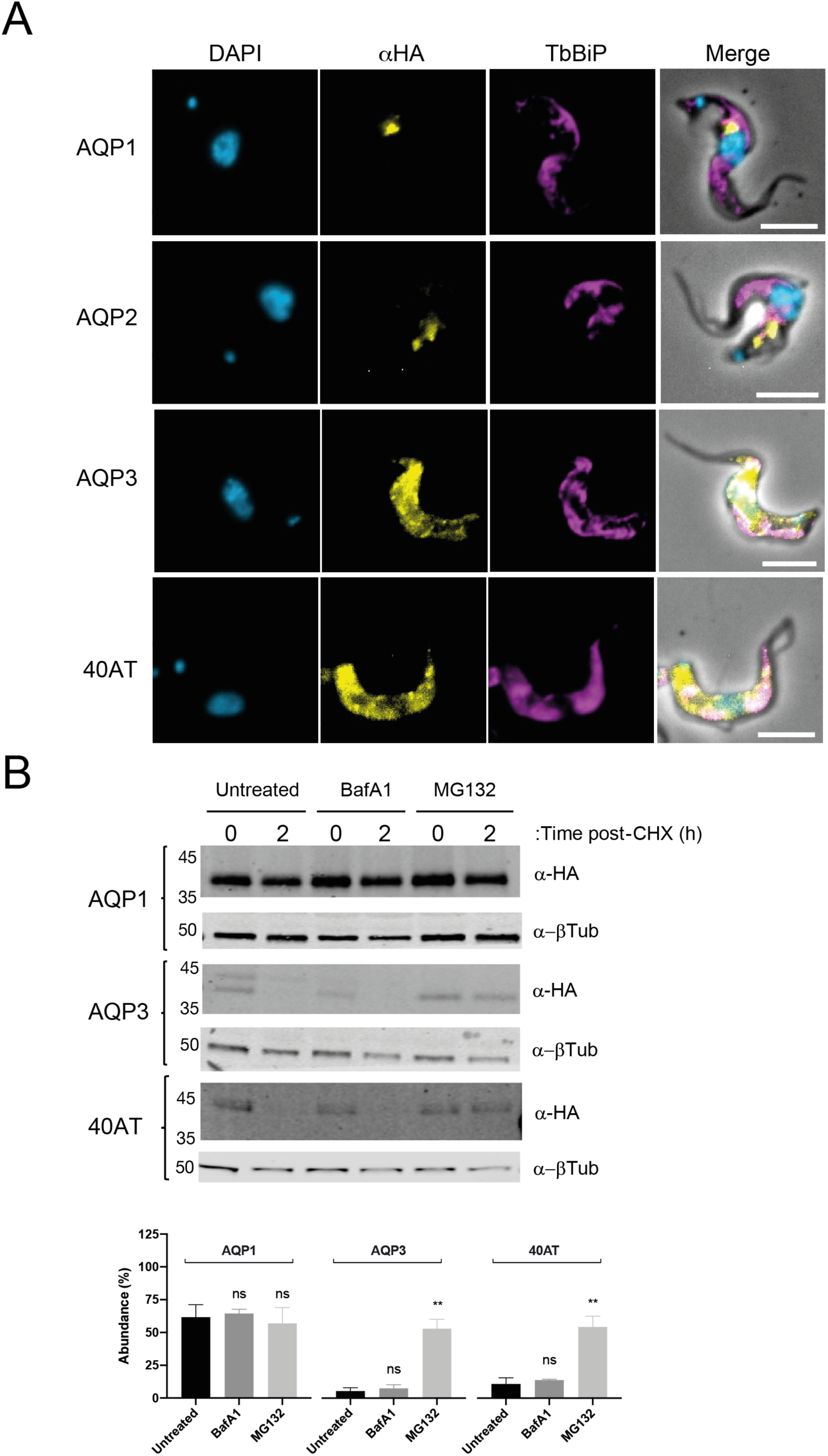
Differential turnover rate of the repertoire of AQPs in the bloodstream form of *T. brucei*. **A)** Cell lines expressing N-terminal HA-tagged TbAQP1, TbAQP2, TbAQP3, field-isolate chimeric AQP2/3 (40AT) or a single TMD mutant (AQP2^TMD4^) (Alexa Fluor 488; yellow) co-stained with the endoplasmic reticulum marker anti-BiP (magenta). DAPI (cyan) was used to label the nucleus and the kinetoplast. Scale bars, 5 μm. **B) Upper panel**; Representative western blot (n = 3 independent replicates) of protein turnover monitored by cycloheximide (CHX) treatment followed by pulse-chase assay. Cells were either untreated or treated with 100 nM of Bafilomycin A1 (BafA1) or 25 μM of MG132 for 1 h prior to harvest. Cells were harvested at 0 hours and 2 h post-CHX treatment and analysed by immunoblotting. **Lowe panel;** Protein quantification representing the mean ± standard deviation of three independent experiments (n = 3 independent replicates). Statistical analysis was conducted using the signal from untreated cells at 2 h post-CHX treatment as reference group. * *p*<0.01, ** *p*<0.001, ns = not significant, using a *t*-test.

**Table 2.** Summary of the impact of chimerisation on TbAQP2 localisation and function.

| **Protein** | **Sequence source** | **Localisation** | **Half-life (h)** | **EC<sub>50</sub> pentamidine (nM)<sup>1</sup>** | **EC<sub>50</sub> SHAM (μM)<sup>1</sup>** | **EC<sub>50</sub> SHAM + 5 mM glycerol (μM)<sup>1</sup>** |
| --- | --- | --- | --- | --- | --- | --- |
| **TbAQP1** | Wild type | Flagellar base | 4.34 | 10.08 | 3.02 | 1.45 |
| **TbAQP2** | Wild type | Flagellar pocket | 4.83 | 3.29 | 1.96 | 1.16 |
| **TbAQP3** | Wild type | Plasma membrane | 1.15 | 10.23 | 1.28 | 1.14 |
| **40AT** | Chimera | Plasma membrane | 1.64 | 10.34 | 1.39 | 1.12 |
| **AQP2<sup>TMD4</sup>** | Chimera | Plasma membrane | <1h | 10.24 | 0.74 | 0.64 |

<sup>1</sup>Estimated EC<sub>50</sub> values from cells induced with tetracycline, 1 μg/ml for 24h.

The localisation of TbAQP3 and the chimeric AQP2-3 proteins is reminiscent of the subcellular localization observed in the lysine-to-arginine TbAQP2 mutants. Based on these observations we hypothesised that these constructs are likely to be retained in the ER, at least to a level comparable to that of the lysine-to-arginine TbAQP2 mutants. We observed that whereas TbAQP1 and TbAQP2 show poor co-localisation with TbBiP, the signal of TbAQP3 and 40AT partly co-localised with this ER marker, indicating some degree of retention within this organelle (**Fig 8A**). Moreover, TbAQP3 and 40AT turnover was faster than TbAQP1 and TbAQP2 and was significantly impaired in the presence MG132 but not bafilomycin A1(**Fig 8B**), indicating that these constructs are retained and degraded in the ER and not in the lysosome, as observed for TbAQP2.

## Discussion

Aquaporins are present throughout prokaryotes and eukaryotes [15–17], and have conserved topology and quaternary structure. AQPs form homotetrameric complexes to transport water and low molecular weight solutes [15–17]. Independent expansions of AQP paralogues have served to diversify function and in mammals and *Leishmania major* both ubiquitylation and phosphorylation are important in modulating turnover and hence activity [61–66]. Significantly, the three trypanosome AQP paralogs derive from a single ancestral gene shared with *Leishmania spp.*, and thus relative functions of paralogs are likely differentially distributed between major lineages. Despite clear clinical importance, little is known concerning AQP trafficking and higher order assembly in trypanosomes and specifically the impact of mutations on these properties. We found that TbAQP2 assembles into high molecular weight complexes that potentially resemble the quasi-arrays described for HsAQP4 [67–71]. Oligomerization correlates with bidirectional glycerol flow but also pentamidine sensitivity as C-terminal tagged forms form oligomers with low efficiency and have poor transport activity. Furthermore, TbAQP2 is ubiquitylated and targeted to the lysosome, similar to mammalian AQPs [62].

Pentamidine is thought to be taken up *via* translocation by and/or endocytosis of TbAQP2 [22,56]. If endocytosis were the sole route and assuming that lysosomal delivery is required, a faster turnover rate than ∼1 h would be necessary to achieve the intracellular pentamidine levels observed, i.e. ∼16 pmol pentamidine/10^7^ cells per hour [72]. Neither pentamidine nor melarsoprol sensitivity requires an obvious lysosomal transporter, suggesting that channel-mediated transport is required, regardless of any endocytic contribution. However, the intrinsic instability of several tagged TbAQP2 mutants precluded detailed dissection of uptake pathways as all of the lysine to arginine mutations led to ER retention [59,60]. As specific mutation of selectivity pore residues does not alter localisation to the flagellar pocket, residues elsewhere are more important for efficient folding.

All *T. brucei* AQP paralogs are predicted as topologically similar, but nevertheless possess distinct properties and subcellular localisations [13,18–23,73]. TbAQP2 is essential for pentamidine and melarsoprol uptake [13], while TbAQP3 is associated with susceptibility to antimonial compounds including sodium stibogluconate [71], indicating transport specificity. Trypanosomes from patients following melarsoprol treatment failure possess a mutated AQP2/3 locus [20,28,29,56,74], with single nucleotide polymorphisms, AQP2 deletions and several fusions replacing TbAQP2 TMD4, 5 or 6 with TbAQP3 sequences, in most cases without impacting the NPA/NPA and WGYR selectivity pore motifs [29,31,32]. Several chimeras have aberrant subcellular localisations [20,28,29,56,74], indicating that the selectivity filters is comparatively unimportant for targeting. Consistent with this is that both TbAQP1 and TbAQP2 assemble into higher order complexes but TbAQP3 apparently does so less efficiently. Similarly, TbAQP1 and TbA have a long half-life (t_1/2_ >4 h) and restricted subcellular localisation around the flagellar pocket, whereas TbAQP3 is comparatively short-lived (t_1/2_ ∼1 h) and localises mainly to the cell body surface, suggesting a connection between oligomerisation, stability, subcellular localisation and transport activity [67,69–71]. Furthermore, replacement of TMD4 or 6 in TbAQP2 by TbAQP3 sequences (we were unable to generate TMD5 chimeras), as observed in clinically relevant chimeric AQP2-3, led to impaired oligomerisation, ER-retention and decreased stability, strengthening the correlation between oligomerisation, localisation and function.

In summary, we propose that pentamidine uptake depends upon the structural organisation of TbAQP2 and that channel activity is essential. Furthermore, TbAQP2 is highly sensitive to mutation and/or chimerisation, which results in failure to correctly fold and ER-retention. This mechanism most likely accounts for many instances of clinically observed pentamidine and melarsoprol resistance.

## Acknowledgements

This work was supported by grants from the Wellcome Trust (204697/Z/16/Z to MCF and 1000320/Z/12/Z to DH) and the Medical Research Council (MR/P009018/1 to MCF). We are grateful to Dave Ng for assistance with sample preparation for imaging and to Martin Zoltner and Ricardo Canavate del Pino for helpful discussions during the preparation of this manuscript. We are also grateful to Pascal Mäser (Swiss Tropical and Public Health Institute) for providing the 40AT chimera.

## Supplementary material for

### Supplementary figure legends

**Figure S1.**
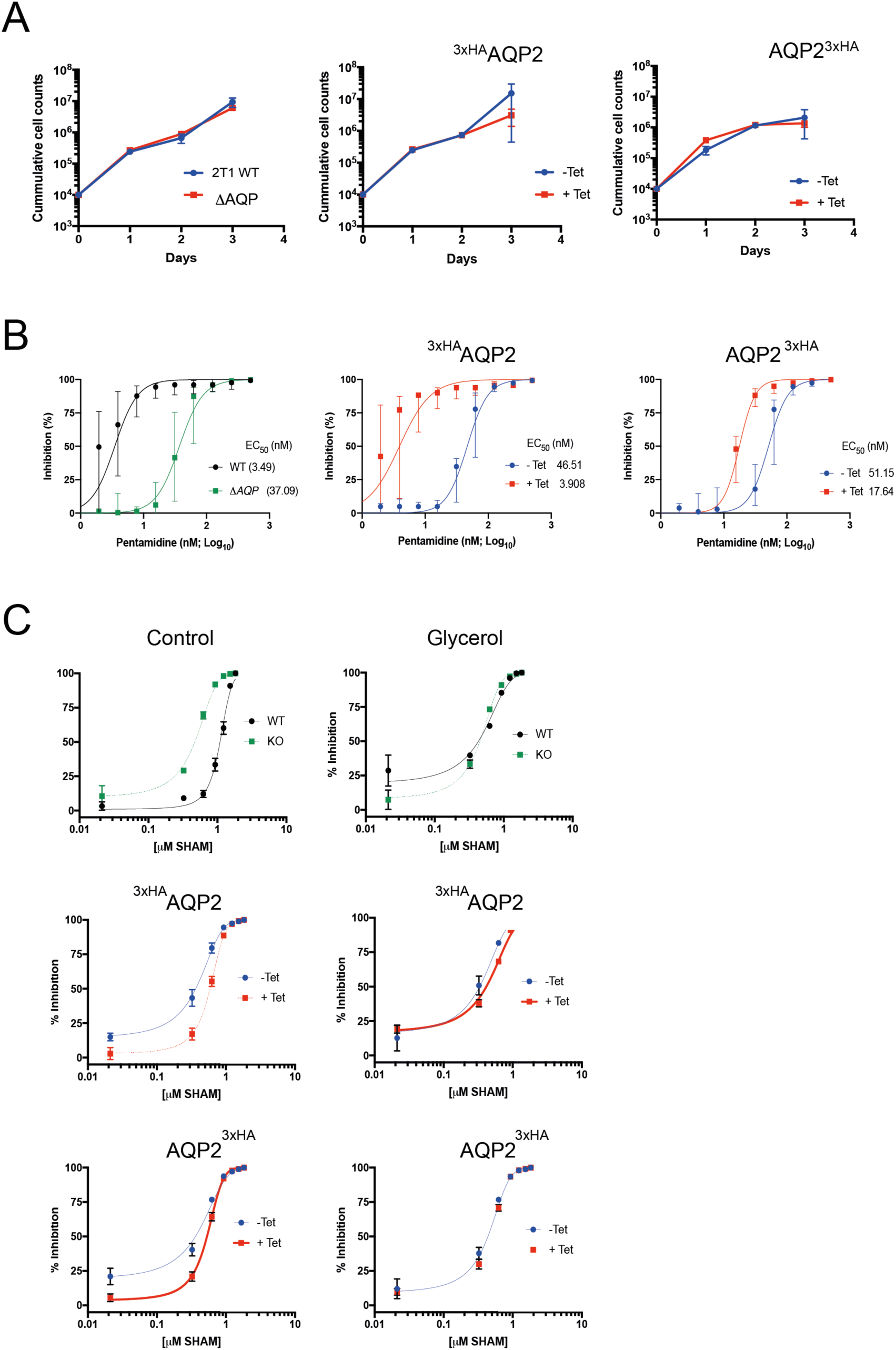
Characterisation of *T. brucei* 2T1 cell lines expressing N- or C-terminal tagged TbAQP2. **A)** Proliferation was estimated over a period of four days *in vitro* in the presence or absence of tetracycline for ^3xHA^AQP2 (middle panel) or AQP2^3xHA^ (right panel) cell lines. *T. brucei* 2T1 wild type *aqp-*null cell lines (left panel) were included as the parental strain. **B)** Dose-response curves for pentamidine from *T. brucei* 2T1 wild type of *aqp-*null cell lines (left panel), ^3xHA^AQP2 (middle panel), and AQP2^3xHA^ (right panel). **C)** Dose-response curves for SHAM (left panels) or SHAM plus 5 mM glycerol (right panels), with the same lines as in panels A and B.

**Figure S2.**
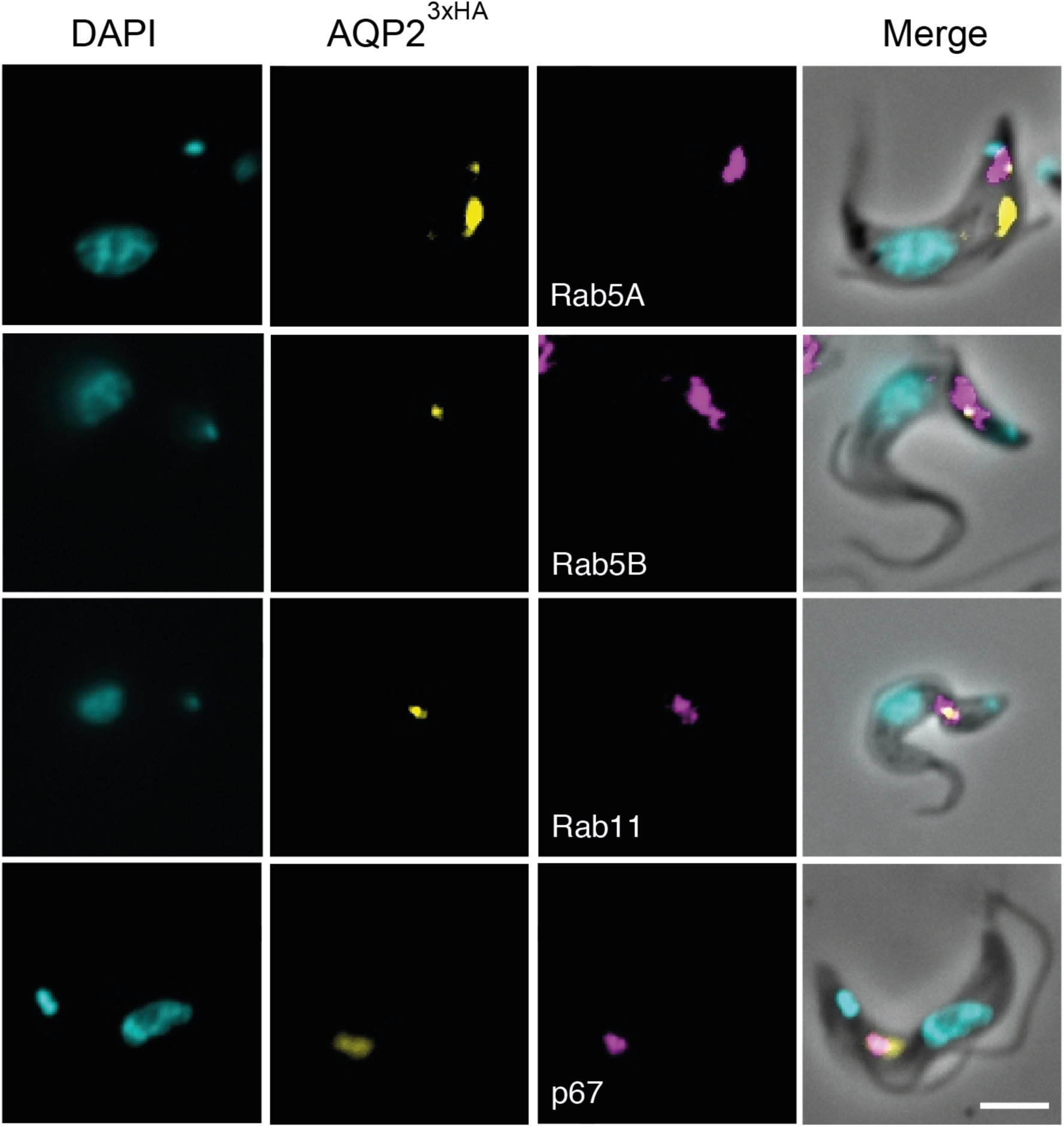
3xHA C-terminally tagged AQP2 transits through endosomes and is delivered into the lysosome. Cell lines expressing a tetracycline-regulated copy of AQP2^3xHA^ (Alexa Fluor 488; yellow) were co-stained with anti-TbRab5a and anti-TbRab5b (early endosomes), anti-TbRab11 (recycling endosomes) and anti-p67 (lysosome). All endosomal and lysosomal markers were labelled with secondary antibodies coupled to Alexa Fluor 568 (magenta). DAPI (cyan) was used to label the nucleus and kinetoplast. Scale bars, 5 μm.

**Figure S3.**
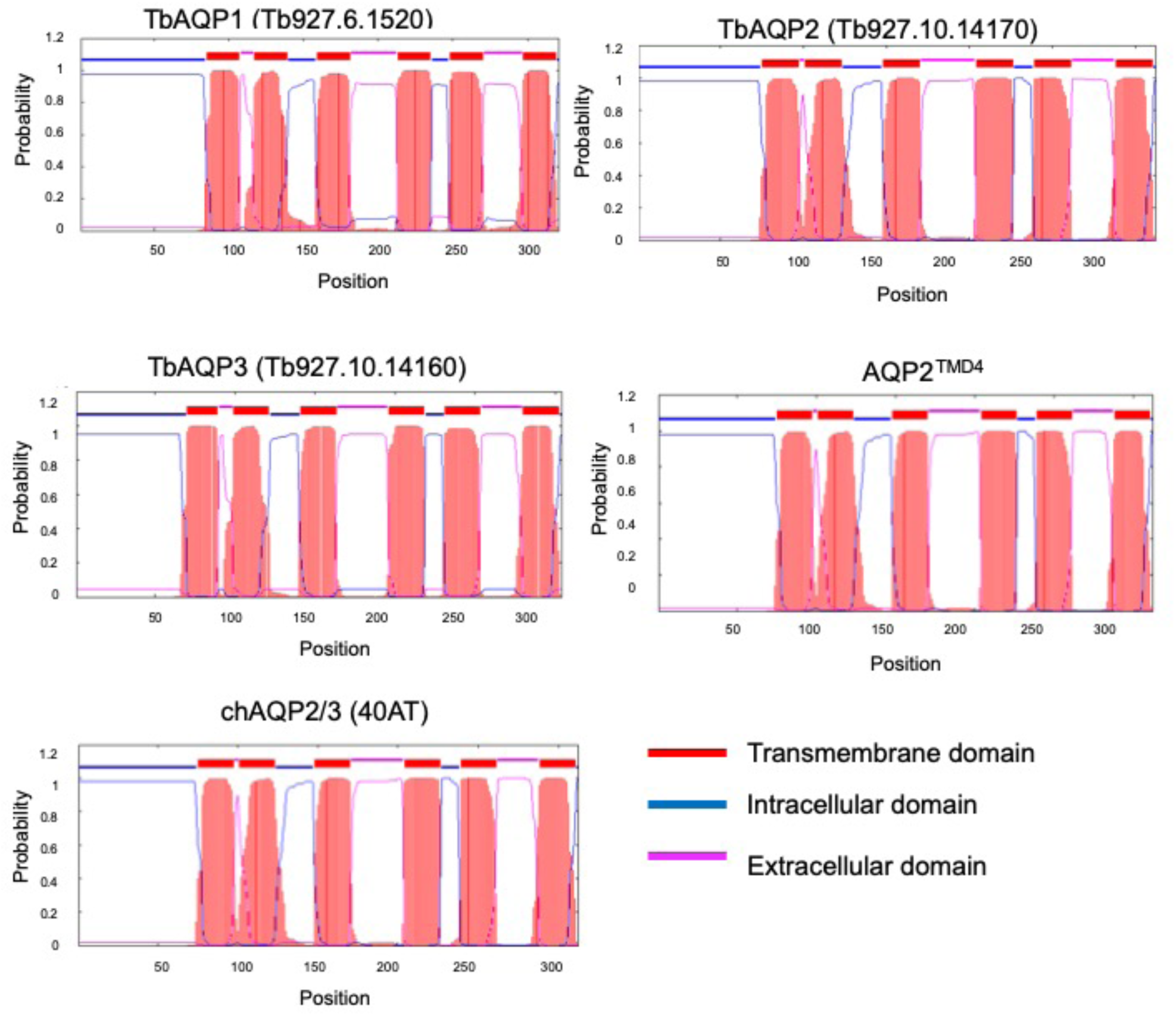
Topological predictions of kinetoplastid aquaglyceroporins. TbAQP1, TbAQP2, TbAQP3, a field-isolated chimera AQP2/3 (40AT) and a single TMD mutant (AQP2^TMD4^) are predicted to have both N- and C-termini facing the cytoplasm and six TMD. Predictions were generated using TMHMM (v2.0) [55].

**Figure S4.**
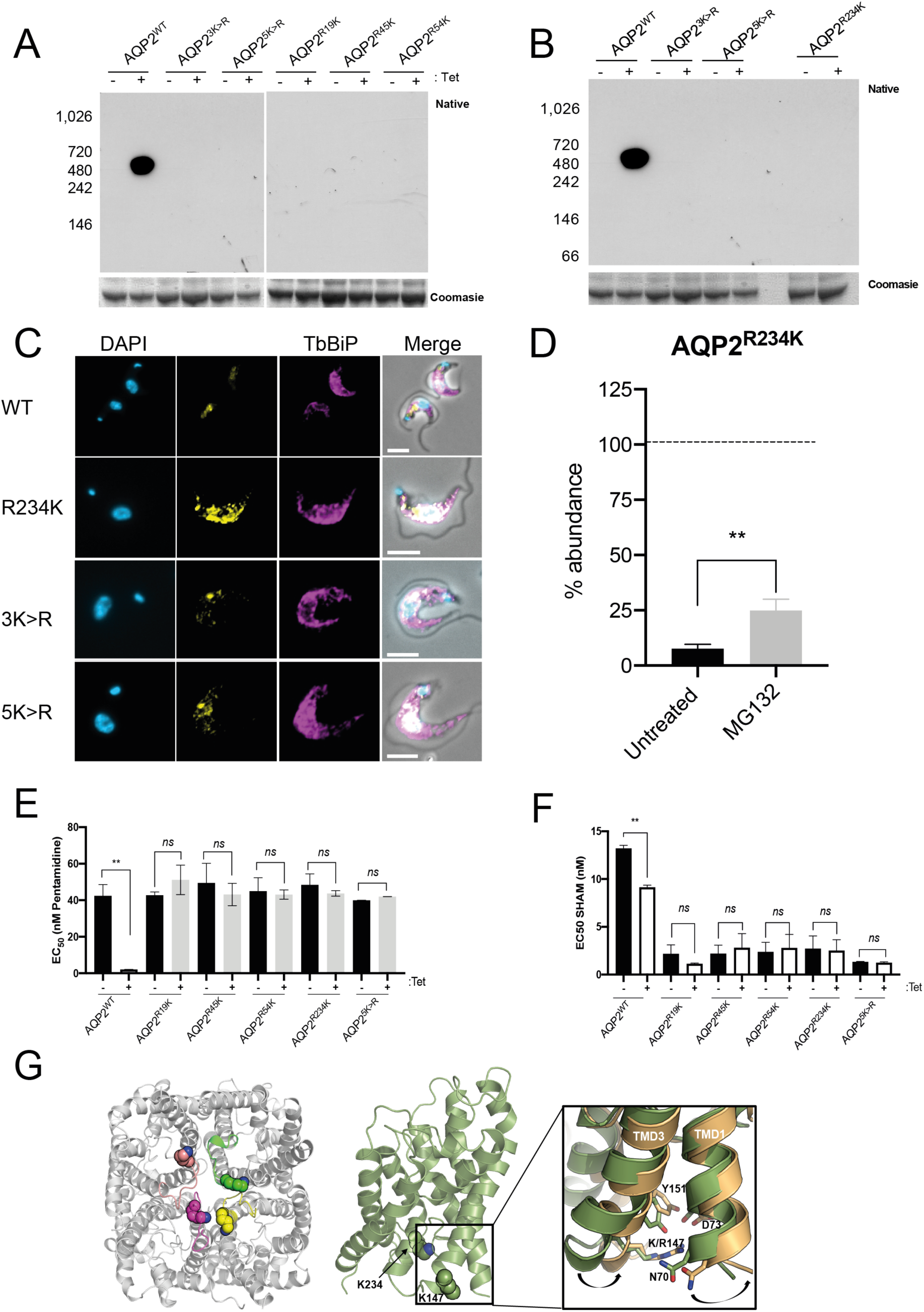
Characterisation of *T. brucei* 2T1 cell lines expressing N-terminal tagged TbAQP2^R234K^ mutant. **A)** Blue native-PAGE immunoblot of total cell lysates expressing either TbAQP2 (AQP2^WT^), N-terminal lysine-to-arginine mutant (AQP2^3K>R^), all lysine-to-arginine mutant (AQP2^5K>R^) and individual arginine-to-lysine mutants (AQP2^R19K^, AQP2^R45K^, and AQP2^R54K^). Coomassie blue staining of the same lysates was used as loading control. **B)** Blue native-PAGE immunoblot of total cell lysates expressing the constructs in (A). Coomassie blue of the same fractions was used as loading control. **C)** Cell lines expressing a tetracycline-regulated copy of wild type TbAQP2 (AQP2^WT^), N-terminal lysine-to-arginine mutant (AQP2^3K>R^), all lysine-to-arginine mutant (AQP2^5K>R^) or individual arginine-to-lysine mutants (AQP2^R234K^) (Alexa Fluor 488; yellow) were co-stained with anti-TbBiP (endoplasmic reticulum) coupled to Alexa Fluor 568 (magenta). DAPI (cyan) was used to label the nucleus and the kinetoplast. Scale bars 5 μm. **D)** Protein turnover was monitored by cycloheximide (CHX) treatment followed by chase and western blotting. Prior to treatment, cells were either untreated or treated with 25 μM MG132 for 1 hour. Cells were harvested at 0 and 2 hours post-CHX treatment and cell lysates analyses by western immunoblotting. Quantification represents mean ± standard deviation (n = 3 independent replicates), and dotted line represents protein abundance at time 0h. Statistical analysis was conducted using untreated cells at 2 hours as reference group. ** *p*<0.001, *ns* = not significant, using a *t*-test. EC_50_ values for pentamidine **(E)** and salicylhydroxamic acid (SHAM) **(F)** with or without 5 mM glycerol following expression of AQP2^R234K^. Statistical analysis was conducted using untreated cells as reference group. ** *p*<0.001, ns = not significant, using a *t* test. Note that this is an extended version of Figure 6A and 6B, and the full panel included for comparison. **G) Left panel**; View from the cytoplasmic face The TMD4-TMD5 loops in each monomer are highlighted. K234 is shown as spheres. **Right panel**; Structural overview of *T. brucei* AQP2 homology model. K147 and K234 are shown as spheres. The expanded view of the conformational change observed during TMD simulations on TMD1 and TMD3 as a result of the K147R mutation. Wild type TbAQP2 is shown in green. TbAQP2 displaying the K147R and K234R mutations is shown in light orange.

**Figure S5.**
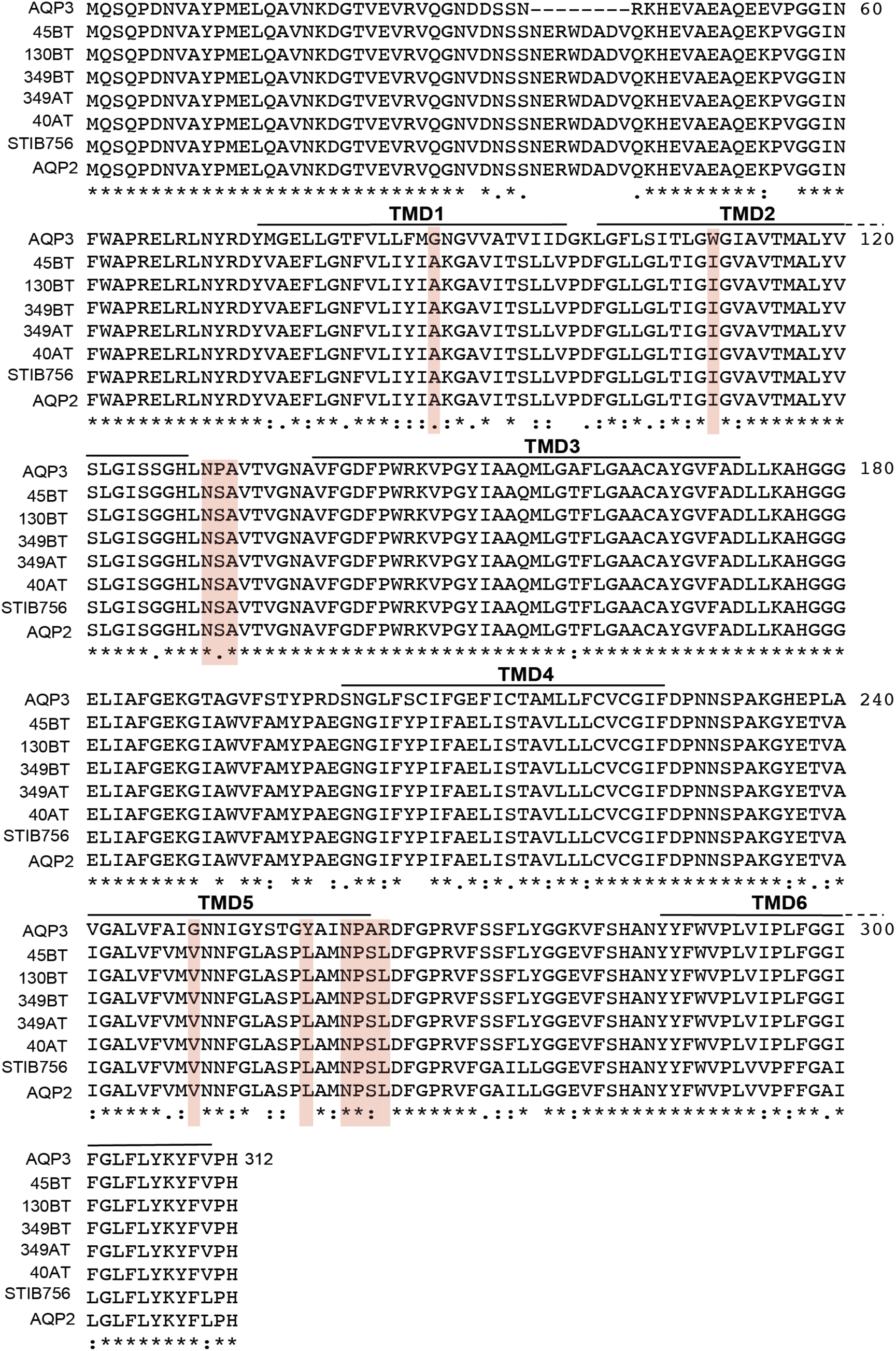
Protein sequence alignment of wild-type TbAQP2, TbAQP3, and the chimeric AQP2-3 detected in relapsing sleeping sickness patients from the Democratic Republic of Congo. The sequence alignment was conducted using Jalview and MUSCLE for multiple sequence alignment. The NPA/NPS selectivity filter is indicated with red boxes, and predicted *trans*-membrane domain (TMDs) spans are also indicated.

**Figure S6.**
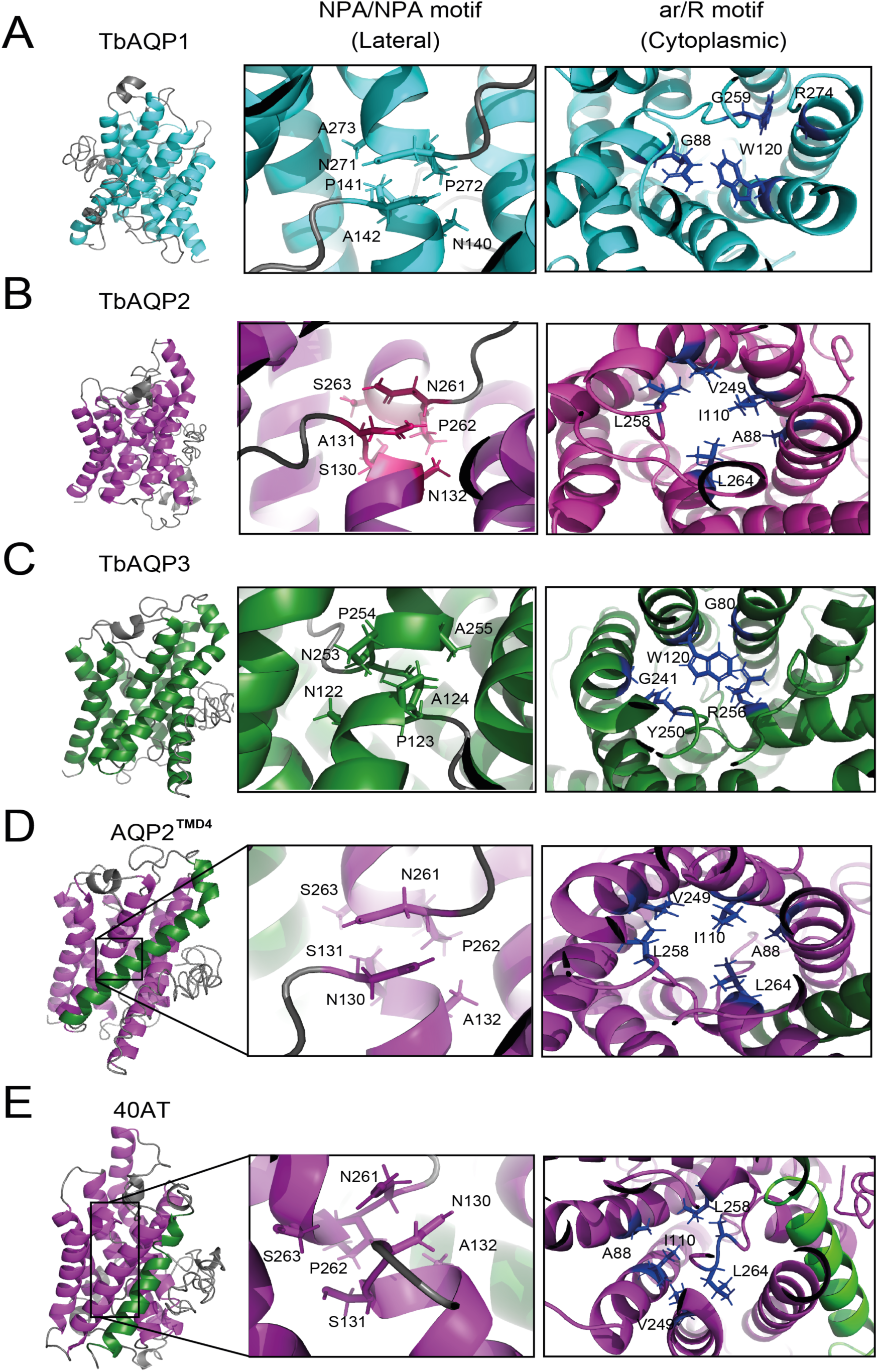
Structural details of TbAQP2 chimerisation and selectivity filters. 3D structural predictions of N-terminal tagged TbAQP1 (cyan), TbAQP2 (magenta), TbAQP3 (green) and the chimeric TbAQP2/3 40AT and AQP2^TMD4^, with the corresponding domain swap colour-coded. Structures were calculated with i-Tasser. The 3xHA-tag is omitted for simplicity. Details of the selectivity pore for each of these proteins (12Å) are also included. For TbAQP2, AQP2^TMD4^ and 40AT constructs, the selectivity filter is composed of the “NSA/NPS” motif, whereas TbAQP1 and TbAQP3 contain a “NPA/NPA” motif.

